# Transcription condensates are promoter hubs that enhance transcriptional bursts

**DOI:** 10.64898/2026.05.24.727549

**Authors:** Ganesh Pandey, Jonah Galeota-Sprung, Alisha Budhathoki, Filmon Medhanie, Jan-Hendrik Spille

**Affiliations:** Department of Physics, University of Illinois Chicago, Chicago, IL, USA; Department of Biomedical Engineering, University of Illinois Chicago, Chicago, IL, USA; NSF-Simons National Institute for Theory and Mathematics in Biology, Chicago, IL, USA

**Keywords:** chromatin, histone modifications, transcription, biomolecular condensates, super-resolution microscopy

## Abstract

Transcription of eukaryotic genes by RNA Polymerase II occurs in temporal bursts and spatial clusters. It is regulated by dozens of transcription factor and coactivator proteins and guided by epigenetic histone marks. Colocalization of transcription machinery in dense foci suggests that cooperative effects orchestrate the process. Factory or condensate models provide a framework for the spatial assembly of the transcription machinery at highly active chromatin loci. But conventional methods lack the resolution to determine how chromatin regulatory elements interact with spatial clusters of the transcription machinery, and whether chromatin structural features modulate functional output. Here, we use super-resolution microscopy to elucidate nanoscale organization of regulatory chromatin at Pol II clusters across scales. We find that Pol II clusters exist on a continuous spectrum of sizes and represent promoter chromatin hubs. We uncover a layered organization of regulatory chromatin, where Pol II clusters form at H3K27ac and H3K4me3-rich domains while H3K4me1 positions peripherally at the surface of large Pol II clusters. Perturbation experiments are consistent with a model in which cohesin loop extrusion forms the active chromatin scaffold underlying transcription assemblies while condensate-driven interactions play only a minor role in genome organization at these sites. Importantly, the number and size of transcriptional burst size increases with Pol II cluster size, revealing directly the cooperative benefits of transcription organization in promoter hubs and a functional consequence of local chromatin structure.

## INTRODUCTION

Transcription of eukaryotic genes is coordinated in space and time. Rather than producing a steady stream of transcripts, productive elongation frequently occurs in transcriptional bursts during which trains of polymerases produce multiple transcripts from the same gene ^1^. Early electron microscopy observations of spatially clustered polymerases led to the concept of transcription factories: static assemblies of polymerases that reel in genes to be transcribed instead of reassembling all parts of the machinery on individual genes ^2,3^. These structures proved elusive in later live cell experiments ^4^, but more recent studies making use of endogenous labeling techniques have uncovered both transient ^5,6^ and stable spatial clusters ^7^ of RNA Polymerase II (Pol II) under unsynchronized and stimulated conditions ^8–11^. Evidence for frequent exchange of constituent proteins ^7,12^ rather than static assembly in structural “factories” is consistent with the notion of transcription condensates ^7,12,13^, although our own single molecule studies indicate that Pol II recruitment to spatial clusters reflects DNA binding rather than phase separation mediated by the Pol II C-terminal domain (CTD) ^14^.

Both stable transcription condensates and transient clusters reflect sites of active transcription and have been shown to modulate transcriptional bursts of individual genes: transient Pol II cluster lifetime determines the number of nascent beta actin mRNA transcripts upon serum stimulation ^8^ and Sox2 transcriptional burst amplitude increases in proximity to condensates ^15^.

However, structural details of chromatin organization at these sites remain poorly resolved. Early electron microscopy data indicates that transcribing polymerases decorate chromatin around a protein core ^16^, while assays using light-activated synthetic condensates suggest that condensates tend to form in regions of low chromatin density ^17,18^. At the same time, condensate forming proteins bound to chromatin can lead to coalescence of decorated elements ^19^. Post-translational histone modifications encode epigenetic information that guides chromatin recruitment of endogenous transcription regulators. Among them, histone 3 (H3), lysine 27 acetylation (H3K27ac) is considered an active chromatin mark that is associated with both promoter and enhancer regulatory chromatin ^20^. This modification has been associated with transcription condensates that accumulate the Mediator complex and the coactivator BRD4 alongside Pol II in dense foci ^7,12^. BRD4 has two bromodomains that allow it to bind to acetylated chromatin ^12,21^. Synthetic multi-bromodomain proteins cause H3K27ac nucleosome arrays to de-mix into micro-phases that are distinct from adjacent unmodified nucleosome arrays condensates ^22^. In zebrafish, H3K27ac chromatin is enriched in large clusters of initiating Pol II ^23^. These findings suggest that BRD4 links active chromatin structures to Pol II clusters, including condensates.

While H3K27ac is a marker for active chromatin in general, H3K4 methylation patterns are more specific for different types of regulatory elements: mono-methylation of H3 lysine 4 (H3K4me1) marks poised enhancers, while tri-methylation (H3K4me3) is found specifically at transcription start sites of active promoters ^20,24^. To the best of our knowledge, the relationship between these more specific marks of chromatin regulatory elements and Pol II clusters has not been investigated.

A universal approach to determine which chromatin loci associate with condensates is lacking – see the accompanying submission by Bogdanović et al. [*Bogdanovi*ć *et al.: Large RNA polymerase II condensates are promoter-centric assemblies.* Reference to be inserted]. In lieu of systematic approaches, prior studies have referred to the strongest peaks in ChIP-seq data to guess candidate genomic loci that may colocalize with transcription condensates. This approach typically yields highly transcribed super enhancer-controlled genes, which indeed colocalize with Pol II clusters to a reasonably high degree ^7,12,23,25^. A severe limitation in our understanding is that sequence-specific chromatin labeling methods in these assays typically cover 10-100kb in linear sequence space and therefore lack the resolution to determine whether specific regulatory elements within the labeled sequences drive the association. Moreover, this approach is too laborious to screen a large panel of genomic loci and remains blind to colocalization of unlabeled interaction partners. Few studies assess colocalization of multiple genomic elements in this context ^26^. Sequencing-based chromatin structure assessment with single cell resolution on the other end of the spectrum provides this information but lacks the protein-level context ^27^.

To overcome these limitations and determine which types of chromatin elements are associated with Pol II clusters and condensates, we conducted two-color, 3D super-resolution microscopy of Pol II isoforms and epigenetic histone marks. Our results show that spatial clustering of Pol II can be explained by the distribution of the Serine 5-phosphorylated, initiating state. Nanoscale spatial organization of regulatory chromatin signatures indicates that Pol II clusters reflect promoter hubs. Transcription burst size in these hubs increases broadly in a manner that depends on the presence of dense Pol II clusters reflecting transcription condensates. Chromatin hubs are mostly resistant to transcription perturbations but enriched in cohesin, leading us to propose a role for DNA loop extrusion in establishing chromatin structures that recruit clustered transcription machinery.

## RESULTS

### Pol II is organized in spatial clusters

Pol II in the cell nucleus is excluded from nucleoli but otherwise appears largely homogeneously distributed throughout the cell nucleus at conventional microscopy resolution (**Fig. 1a,b**). This is expected since at micromolar concentration ^4,28^, individual molecules are separated by approximately 100nm – far less than the resolution limit of conventional optical microscopy. Super-resolution microscopy uncovers a more nuanced picture. Reconstructing super-resolved density maps from single molecule localizations with 10-20nm localization precision (**Suppl. Fig. 1a**) reveals nanoscale Pol II clusters (**Fig. 1a**).

**Fig. 1:**
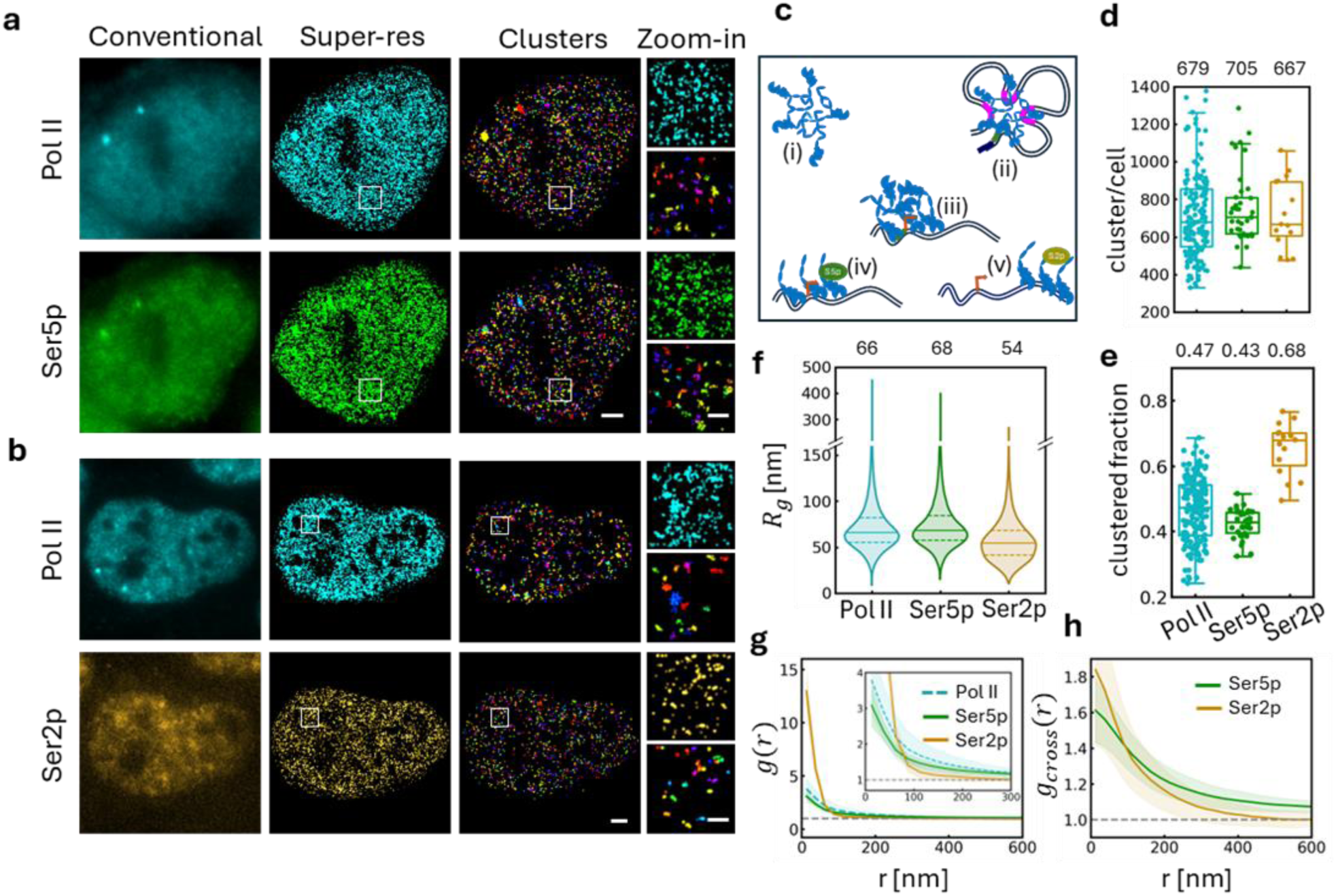
Pol II cluster size distribution reflects initiating Pol II distribution. **A**Immunofluorescence images of global Pol II (top) and initiating Ser5p-Pol II (bottom) acquired by conventional (left) and super-resolution microscopy (middle) as well as nanoscale clusters identified by DBSCAN (right). **b** Same for global Pol II (top) and elongating Ser2p-Pol II (bottom). **c** Several possible mechanisms can explain Pol II clustering, including i) CTD-mediated phase-separation, ii) accumulation in condensates at super-enhancers, iii) transient clustering at promoters, iv) pile-up of promoter-proximal paused Pol II, and v) trains of elongating Pol II in transcriptional bursts. The degree of phosphorylation differs between the scenarios. **d** Number of clusters identified per nucleus in a single optical plane. **e** Clustered fraction per nucleus for the different phospho-isoforms. n=168 (Pol II); 37 (Ser5p); (15) Ser2p. **f** Size distribution (radius of gyration) of clusters. Solid line indicates the median value, dashed lines quartile boundaries. Median values are indicated above each group. **g** Parameter-free pair-correlation analysis reports spatial clustering of Pol II, Ser5p, and Ser2p localizations. Ser5p clustering properties are very similar to general Pol II by all metrics, whereas Ser2p clusters to a higher degree and in smaller foci. **h** Cross pair-correlation function of Pol II and Ser5p (green) or Pol II and Ser2p (orange) localizations. Number of cells: d, f, g: n=168 (Pol II); n=37 (Ser5p); n=15 (Ser2p). h: n=37 (Pol II–Ser5p); n=15 (Pol II–Ser2p). Number of clusters: e: n=121,156 (Pol II); n=28,044 (Ser5p); n=10,864 (Ser2p). Scale bars: **a, b** 2 µm; magnified views 500 nm.

Pol II clusters have been described in various biophysical terms, ranging from transient assemblies ^5^ driven by self-association of the Pol II C-terminal domain (CTD) ^29^, to wetting of active chromatin ^23^, association with condensates at super-enhancers ^7^, stalling at promoter-proximal pause sites ^30^, or elongation in trains of polymerases in transcriptional bursts ^1,23^ (**Fig. 1c**). Throughout most of this manuscript we remain agnostic to formation mechanisms and refer to spatial aggregations of Pol II simply as clusters. The largest Pol II clusters in particular have been characterized further and described as biomolecular condensates ^7,31^. Large condensates are included in our analyses, but we cannot with certainty define criteria or thresholds to distinguish condensates from other types of Pol II clusters. It remains to be determined whether clusters of all sizes reflect the same biophysical assembly mechanisms.

Here, we performed two-color 3D dSTORM super-resolution microscopy on immuno-stained v6.5 mouse embryonic stem cells using a biplane detection approach ^32^ with nanogold fiducials to correct for drift and chromatic aberrations. We then used density-based clustering for applications with noise (DBSCAN) ^33^ to segment clusters from the lists of single molecule localizations. Since DBSCAN output under given parameters depends on localization density, we subsample localizations by random thinning to the same localization density in each nucleus. This allows us to comparatively analyze all nuclei with the same set of DBSCAN parameters. The procedure does not affect global measures of clustering, e.g. parameter-free pair-correlation analysis (**Fig. 1g, Suppl. Fig. 1**). On average, we detect close to 700 individual Pol II clusters in a single focal plane (**Fig. 1d**). The number of clusters scaled linearly with the observed volume per cell, allowing us to extrapolate to an estimated total number of 3500 in the entire nucleus (**Suppl. Fig. 1d,e**) in excellent agreement with prior reports ^28,31^. With our approach, 47% of Pol II localizations obtained with an antibody that recognizes “general Pol” (both unmodified and phosphorylated isoforms) were assigned to clusters (**Fig. 1e**). As measures for cluster size, we calculated the radius of gyration of localizations within a clusters (average size of 66nm) (**Fig. 1f**) and the number of localizations per cluster.

To determine whether clustering properties differ between functional states of Pol II, we repeated the same analysis for phosphorylated Pol II isoforms. Serine 5 phosphorylation (Ser5p) is deposited during the initiation and early stages of transcription and gradually replaced with Serine 2 phosphorylation (Ser2p) in late elongation towards transcript termination ^34^. We used antibodies against either isoform together with a general Pol II label in two-color super-resolution experiments. Clustering assessment by pair-correlation analysis yields very similar amplitude and decay length for Ser5p spatial organization as compared to general Pol II (**Fig. 1g**). DBSCAN cluster analysis is consistent with this observation. Ser5p cluster analysis yields a similar number of clusters per cell, clustered fraction, and size distribution (**Fig. 1d-f**). This is not surprising given that most Pol II is engaged with chromatin and only a small fraction in a freely diffusive state ^14,35^.

We also conducted the same analysis for the late elongating Ser2p isoform (**Fig. 1b**). Overall, super-resolution imaging of Ser2p yielded fewer localizations per cell than general Pol II and Ser5p labeling (**Suppl. Fig. 1c**). This may reflect differences in antibody binding and labeling efficiencies or simply abundance of the epitope. It is not easily possible to disentangle and quantify these effects. We detected a similar number of Ser2p clusters as for Ser5p and general Pol II (**Fig. 1d**), but Ser2p clusters to a higher degree as quantified by pair-correlation analysis (**Fig. 1g**) and DBSCAN cluster metrics (68% clustered fraction, **Fig. 1e**). The pair-correlation function falls of more steeply and at shorter length scales, confirming that Serp2 clusters are smaller than general Pol II and Ser5p clusters (**Fig. 1d,e**).

To quantify the spatial relationship between general Pol II and its phosphorylated isoforms, we computed cross-pair correlation functions. Unlike binary co-localization measures, this approach resolves spatial associations between two targets as a function of length scale. Both Ser5p and Ser2p yield similar profiles in cross-pair correlation analysis (**Fig. 1h**), indicating colocalization of general and phospho-specific Pol II signal on length scales of up to 200nm.

### Active chromatin density is a predictor of Pol II clustering

To directly quantify association of Pol II clusters with active chromatin in single cells, we acquired two-color super-resolution images of general Pol II and the three active chromatin marks H3K27ac, H3K4me1, and H3K4me3. We identified Pol II clusters using DBSCAN (**Fig. 2a**) as described above and evaluated their coordinates in super-resolved maps of active chromatin density (**Fig. 2b**). To quantify active chromatin density, we computed the Gaussian kernel density of histone mark localizations with a bandwidth of 100nm. We then normalized maps for each nucleus to the mean density and quantized into 20 bins on a log2-scale, where 0 corresponds to the average signal throughout the nucleus (**Fig. 2b**). To determine how likely Pol II clusters were to be found in regions of a given chromatin density, we computed the ratio of observed frequency to random expectation. This yields the predictivity score *ρ*^. We plotted *ρ*^ as a function of H3K27ac density for Pol II clusters of different sizes (**Fig. 2c**). Pol II clusters are less likely than randomly expected to form in regions of below average active chromatin density. The predictivity for Pol II cluster location increases sharply for nuclear regions with the highest active chromatin density. This effect is most pronounced for larger Pol II clusters. As previously reported, not all active chromatin regions harbor Pol II clusters ^23^. But regions with the highest active chromatin density are 6-fold more likely than randomly expected to harbor Pol II clusters of >200nm radius. These results establish the propensity of the most active chromatin regions to recruit Pol II clusters of all sizes and in particular large condensates.

**Fig. 2:**
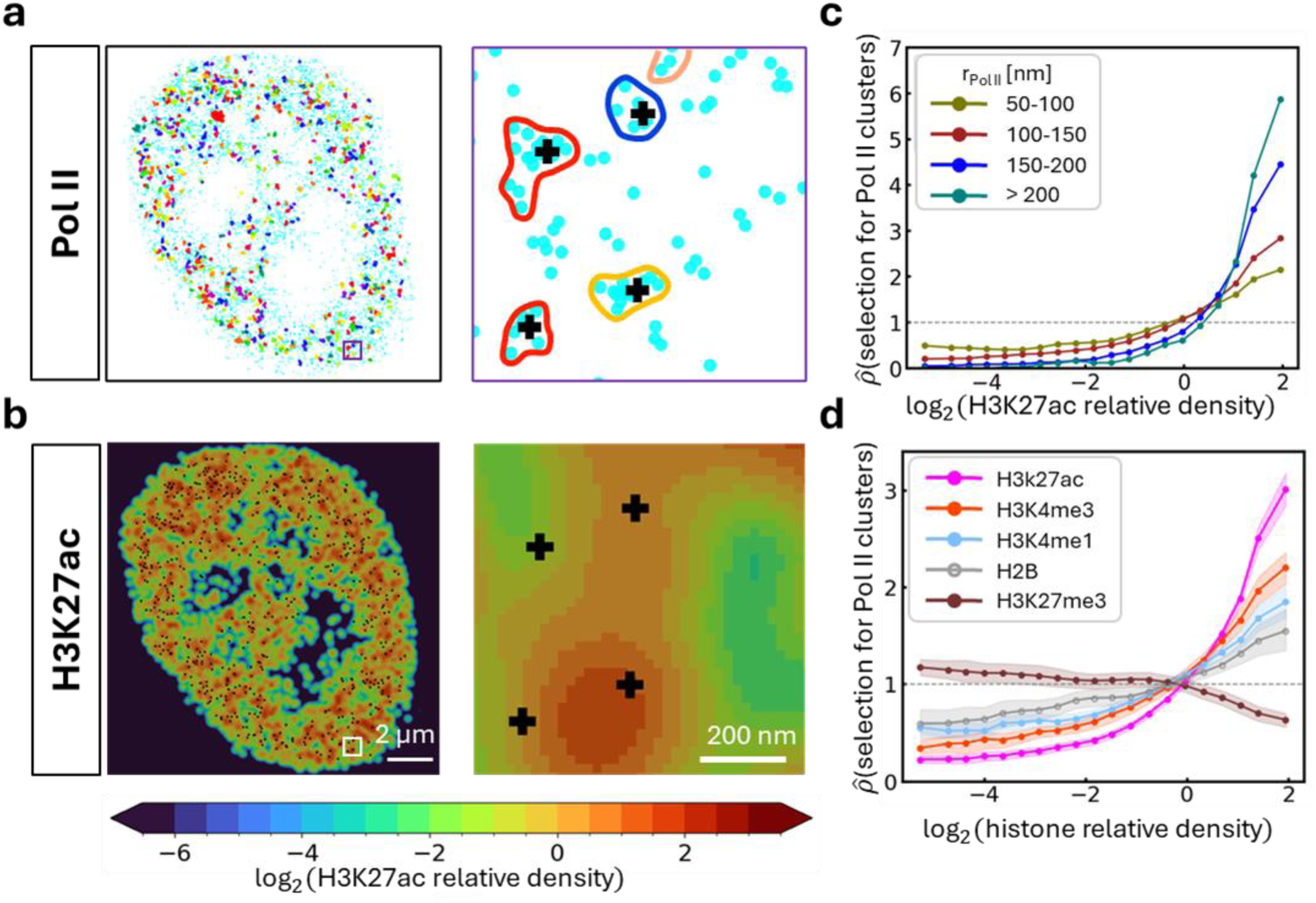
Active chromatin density predicts Pol II cluster location. **a** Scatter plot of 3d dSTORM Pol II localizations (cyan). DBSCAN clusters are highlighted in colors. Right: Magnified view highlighting cluster outlines and centroids (+). **b** Kernel density (bandwidth 100nm) of H3K27ac 3d dSTORM localizations quantized into 20 bins. Colors reflect fold density compared to average in the nucleus. Right: Magnified view of Pol II cluster centroids (+) overlaid on H3K27ac density. **c** *ρ*^ predictivity score for Pol II cluster selection as a function of H3K27ac density stratified by Pol II cluster size (radius of gyration). Highest H3K27ac density regions are most predictive of Pol II clusters. Predictivity increases for larger Pol II clusters. **d** *ρ*^ for Pol II clusters of all sizes for different chromatin marks. H3K27ac and other active chromatin marks (H3K4me1/3) are most predictive of Pol II clusters, high density regions of inactive chromatin (H3K27me3) are depleted of Pol II clusters. Scale bars: **a, b**, full nucleus 2 µm; magnified view 200 nm.

We observe qualitatively similar trends for the H3K4me1 and H3K4me3 marks (**Fig. 2d**). Each of the three active chromatin marks is predictive of Pol II clusters of all sizes, but most predictive for the largest clusters (**Suppl. Fig. 2**). Out of the three active histone marks, H3K27ac was most predictive of Pol II clusters, followed closely by H3K4me3. H3K4me1 predictivity scores were lowest out of the three marks assessed here and peaked at approximately 3-fold enrichment for the largest Pol II clusters (**Suppl. Fig. 2c**). Optimizing predictivity of H3K4me1 maps required a slightly larger kernel density radius than H3K27ac and H3K4me3 density maps (**Suppl. Fig. 2a**). This is somewhat surprising since H3K4me1 is considered an enhancer mark and transcription condensates, i.e. the largest Pol II clusters, are thought to associate with super-enhancers ^12^. H3K4me3 predictivity increases sharply for the largest Pol II clusters (**Suppl. Fig. 2b**). As a negative control we conducted the same analysis for the repressive chromatin mark H3K27me3. As expected, Pol II clusters of all sizes are less likely to be found in regions with high H3K27me3 density (**Fig. 2d, Suppl. Fig. 2e**).

Notably, predictivity extends to H2B density (**Fig. 2d**), which we included as a marker of nucleosomal chromatin regardless of epigenetic state. H2B density correlates mildly with Pol II cluster abundance for clusters of all sizes (**Suppl. Fig. 2d**). Differences in predictivity emerge mostly for the lowest H2B densities. Interestingly, the largest Pol II clusters are least likely to occur in regions of very low H2B density. We also note a small deviation for the largest Pol II clusters (>200nm). Predictivity for this class peaks at an H2B density corresponding to approximately 2-fold enrichment above average but decays towards higher densities of 4-fold above average. This indicates that the largest Pol II clusters in particular preferentially form at a specific chromatin packing density ^36,37^. These observations are in agreement with prior reports that large Pol II clusters overlap with active histone marks in zebrafish ^23^ and contrast with synthetic condensates that preferentially form in regions of low chromatin density ^17^.

### Active chromatin domains extend beyond Pol II clusters

The observation that the predictivity score increased for larger Pol II clusters led us to ask whether clusters of different size associate with distinct chromatin environments. To answer this question, we binned Pol II clusters by size (radius of gyration) and aligned super-resolution localizations in both channels with respect to individual Pol II cluster centroids (**Fig. 3a**). We aggregated localizations from all regions of interest in both channels and normalized localization density to the number of contributing clusters.

**Fig. 3:**
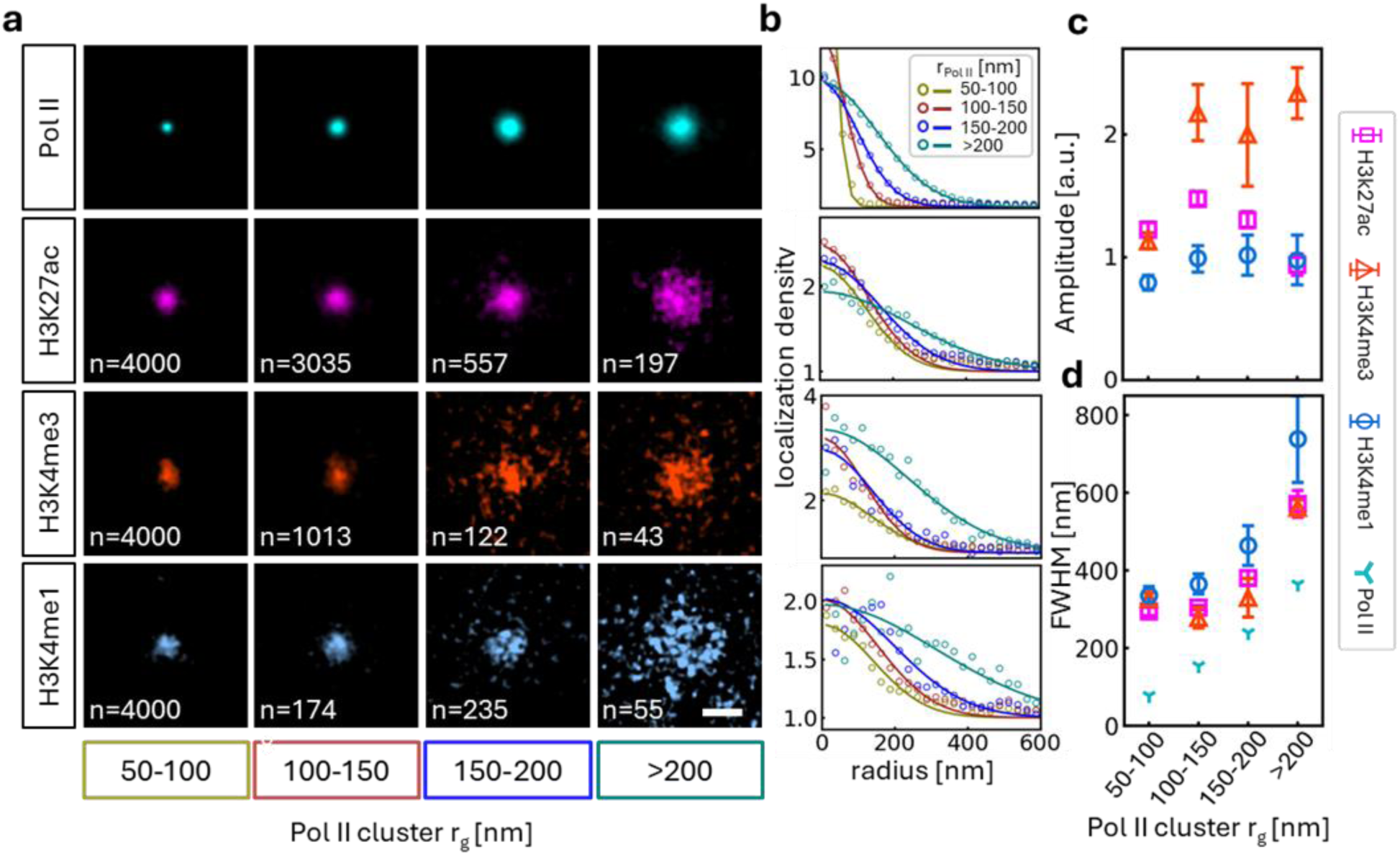
Active chromatin domains extend beyond Pol II clusters. **a** Aggregated two-color super-resolution localizations stratified by radius of gyration of Pol II clusters. Localizations were centered on Pol II cluster centroid coordinates for Pol II (top). Numbers represent the number of Pol II clusters considered for the respective histone mark data. **b** Radial localization density normalized to the number of contributing clusters and background density (mean value at 500nm-1000nm radius). We fitted with a normal distribution to facilitate comparison of **c** localization density amplitude and **d** spread (full width at half maximum, FWHM) of the signal. H3K4me3 density increases notably for larger Pol II clusters. The spread of all active chromatin marks increases with Pol II cluster size and exceeds the spread of Pol II localizations. H3K4me1 is enriched at larger distances than H3K27ac and H3K4me3. Data in **c** and **d** represent bootstrapped mean and standard deviation. Scale bar: a, 200 nm.

In this aggregate analysis, all three active chromatin marks are associated with Pol II clusters of all sizes. To systematically assess the nature of active chromatin enrichment, we quantified chromatin mark density as a function of radial distance from the Pol II cluster centroid. Fitting with a normal distribution yields an amplitude above background (A) and spread (full width of half maximum, FWHM). The amplitude corresponds to the active chromatin concentration at Pol II cluster coordinates, whereas the spread reports how far from cluster centroids active chromatin concentration is elevated (**Fig. 3b**).

The concentration of H3K27ac and H3K4me1 showed only small variation with Pol II cluster size around an amplitude of 1 over background, i.e. twice the average density in the nucleus (**Fig. 3c**). H3K4me3 concentration increased for Pol II clusters >100nm to 3-fold the nuclear average. This analysis is distinct from the one presented in **Fig. 2** since here, we change the viewpoint and focus on coordinates of detected Pol II clusters rather than chromatin spatial organization. The analysis presented here also works with the full localization precision instead of calculating local density with a 100nm radius Gaussian kernel.

Elevated epigenetic signal extends in domains beyond the spread of Pol II localizations at clusters of all sizes (**Fig. 3d**). Even for the smallest Pol II clusters considered (50-100nm), the active chromatin signal spreads over more than 300nm (FWHM). The corresponding metric of Pol II localization yields only 80nm (FWHM). The spread of chromatin signal increases only slightly with Pol II cluster size of up to 200nm. This indicates that Pol II clusters of all sizes are embedded in active chromatin domains of a characteristic size with approximately 300nm FWHM, corresponding to features observed by electron microscopy ^37^. At the largest Pol II clusters, active chromatin spreads over more than 550nm FWHM (H3K27ac and H3K4me3) or up to 740nm FWHM (H3K4me1).

H2B signal as a proxy for nucleosomal chromatin is enriched in particular at smaller Pol II foci, again suggesting that Pol II clusters associate with active chromatin patches rather than forming off chromatin (**Suppl. Fig. 3a**). H2B concentration decreases at larger Pol II clusters but remains elevated over local background. The spread of H2B signal tracks closely with the spread of Pol II itself, supporting the notion that Pol II clusters reflect chromatin structures (**Suppl. Fig. 3b,c**). Cross-pair correlation analysis of Pol II and histone mark localizations sheds more light on the relationship between Pol II clusters and chromatin density. H2B density is elevated at short distances <100nm around Pol II, but depleted at larger distances (**Suppl. Fig. 3e**). This indicates local contraction at sites of Pol II clusters. H3K27me3 inactive chromatin signal is depleted at Pol II clusters of all sizes and most strongly for larger Pol II clusters (**Suppl. Fig. 3**).

These results show that chromatin zones of influence extend well beyond the boundaries of Pol II clusters. Notably, the H3K4me3 promoter mark is most enriched at Pol II clusters in this analysis, whereas the H3K4me1 enhancer mark spreads furthest outside Pol II clusters.

### Promoter marks are interior to large Pol II foci, enhancer marks peripheral

Our chromatin analysis at Pol II clusters has so far focused on aggregate signal around Pol II cluster centroid coordinates. This approach yields intriguing insights but averages out variation in size and shape between individual Pol II clusters. To overcome this limitation, we developed a more sophisticated approach that takes into account the spatial organization of chromatin around individual Pol II clusters. We shifted the vantage point to the surface of segmented Pol II clusters defined by the alpha-hull around each cluster (**Fig. 4a**). Magnified views of representative clusters demonstrate their irregular shapes. We then quantified localization density as a function of orthogonal distance to the cluster surface (**Fig. 4b**). We normalized data to values obtained by repeatedly placing the same Pol II cluster outlines at random coordinates within the same nucleus. The result is a set of radial enrichment profiles relative to the Pol II cluster surface rather than measured from aggregate centroids (**Fig. 4c**).

**Fig. 4:**
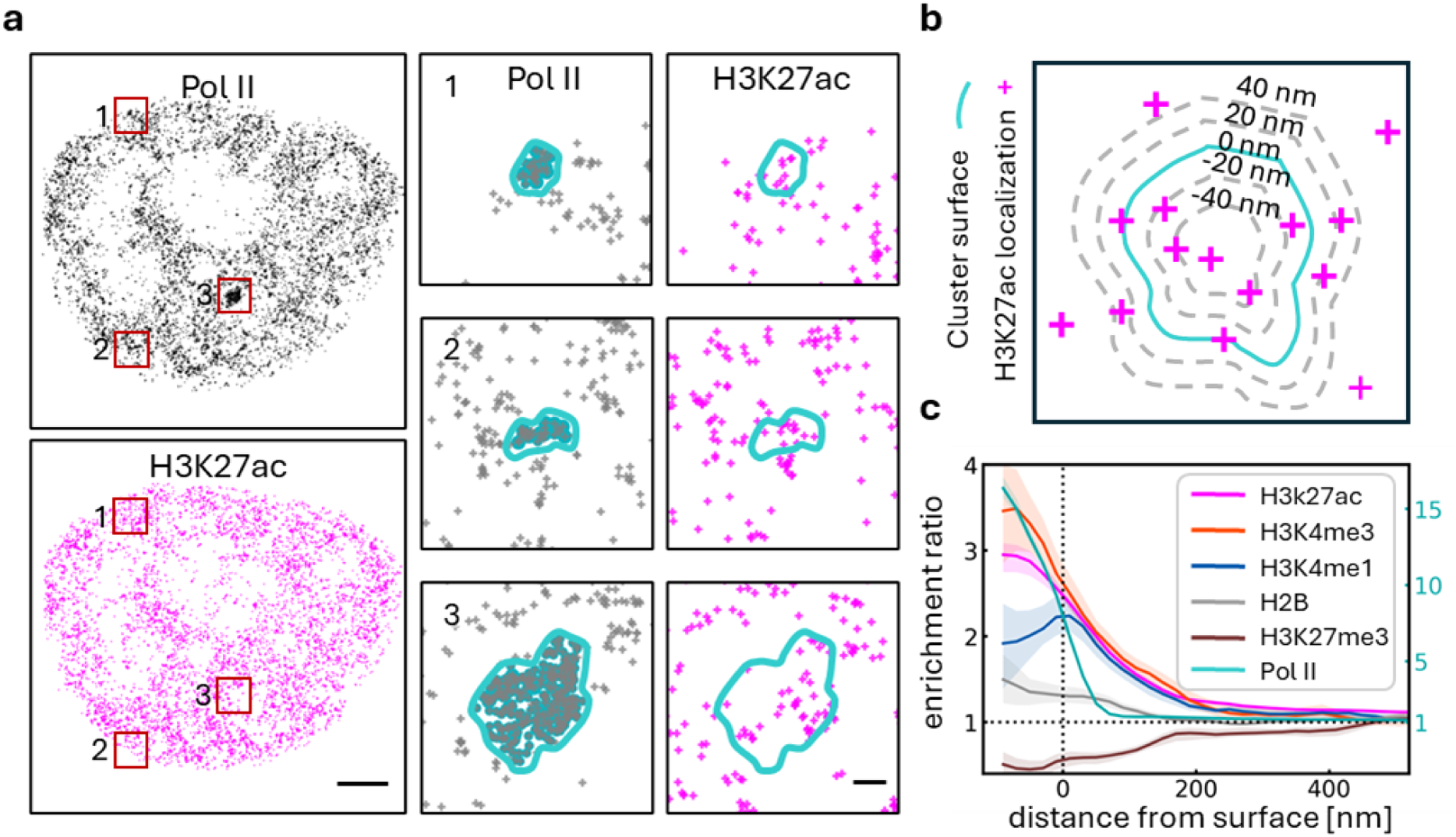
Differential spatial distribution of promoter- and enhancer-marks around Pol II clusters. **a** Single-molecule localizations of Pol II (gray) and H3K27ac (magenta). Left: Full nucleus. Right: Magnified views centered on three representative Pol II clusters of different sizes (gray outlines) overlaid with cluster surfaces (cyan contour) and single H3K27ac localizations (magenta). **b** We measured the orthogonal distance of each histone mark localization relative to the Pol II cluster surface. Gray dashed lines indicate equidistant contours. Distances inside the cluster are negative, outside the cluster positive. **c** Enrichment profile of histone mark localization density around Pol II clusters with radius of gyration ≥150 nm normalized to profiles obtained by repeating the analysis with the same cluster surfaces at randomly chosen coordinates in the same nuclei. H3K27ac and H3K4me3 enrich towards the core of Pol II clusters. H3K4me1 localization density peaks precisely at the Pol II cluster surface. Scale bar: **a** full nucleus 2μm; magnified view 200nm.

We conducted this analysis initially only for the larger Pol II clusters >150nm radius (**Fig. 4c**). Both H3K27ac and H3K4me3 localization density enriches at negative distances from the surface, i.e. towards the Pol II cluster interior. As before, H3K4me3 enrichment is higher than H3K27ac enrichment. In contrast, the H3K4me1 enhancer mark density peaks precisely at the Pol II cluster surface and falls off in both directions. This surprising finding suggests a layered organization of regulatory chromatin and distinguishes the Pol II cluster surface from its interior. The cluster interior consists of a chromatin core that is rich in promoter (H3K4me3) and active enhancer (H3K27ac) marks, while poised enhancers (H3K4me1) are positioned peripheral to this assembly, mirroring the wider spread of this mark (**Fig. 3**).

As an inherent control, we conducted the same analysis on the Pol II channel itself. Pol II localization density falls off sharply at the inferred cluster surface over a range of ±50nm. This demonstrates the precision of our surface segmentation approach. Histone mark density falls off more slowly over distances of up to 200nm from the surface. The observed effect is therefore not a technical artifact of the segmentation.

Core enrichment of H3K27ac and H3K4me3 is conserved at smaller Pol II clusters, whereas H3K4me1 shifts from surface towards core enrichment for smaller clusters (**Suppl. Fig. 4**). We note, however, that defining irregular surfaces and measuring orthogonal distance from them becomes increasingly difficult for small Pol II clusters. Controls for nucleosomal chromatin (H2B) and repressed chromatin (H3K27me3) yield expected results: H2B signal is weakly enriched to the same degree across all cluster sizes whereas H3K27me3 is broadly depleted inside and around Pol II clusters (**Fig. 4c, Suppl. Fig. 4**).

### Active chromatin marks form nanodomains

Super-resolution microscopy also reveals clustered nanodomains of all three epigenetic marks as previously reported ^38^ (**Fig. 5a**). We again used DBSCAN to identify 700-1000 active chromatin nanodomains per nucleus in a single focal plane (corresponding to 3500-5000 per cell) (**Fig. 5b**). H3K4me3 yields the highest number of nanodomains, comparable to the H2B control. H2B nanodomains have previously been described as clutches of ∼10 nucleosomes concentrated in clusters with a radius of tens of nanometers ^39^. This is in good agreement with our quantification.

**Fig. 5:**
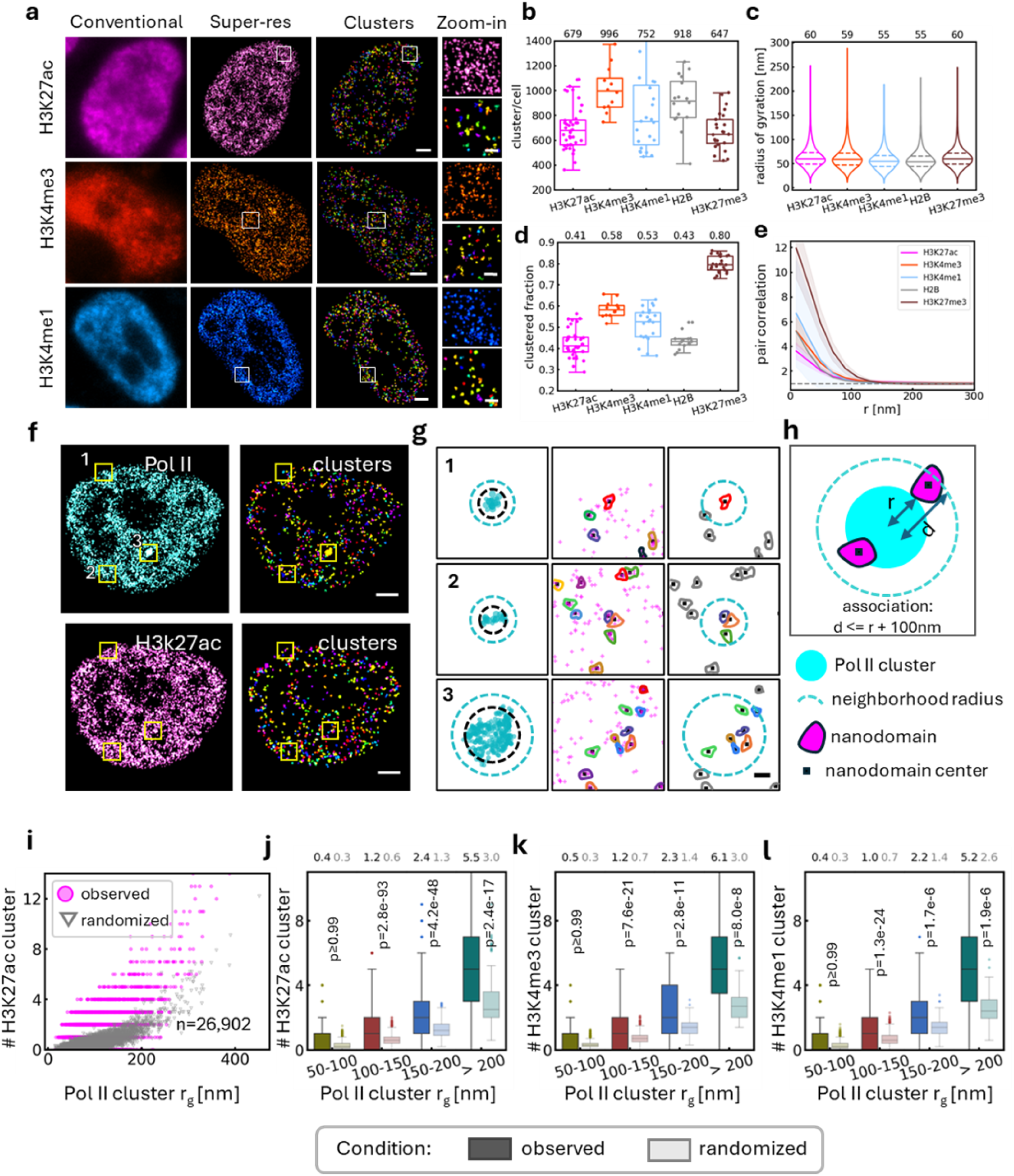
Active chromatin forms clustered nanodomains. **a** Fluorescence microscopy images of active chromatin marks at conventional resolution (left) and in super-resolution (middle). DBSCAN clusters identified from super-resolved images (right) highlight chromatin nanodomains. Magnified views of all localizations (top) and clustered localizations colored by ID (bottom). **B** Number of clusters per focal plane in single nuclei. **c** Size distribution of chromatin nanodomains. **d** Clustered fraction of localizations per nucleus. H3K27ac clusters to the lowest degree. The inactive chromatin mark H3K27me3 shows the highest degree of clustering. H2B clusters to a similar degree as active chromatin marks. **e** Pair-correlation analysis of chromatin mark localizations indicates clustering at short distances <100nm, consistent with DBSCAN cluster sizes. **f** Rendered super-resolution images of Pol II and H3K27ac in the same nucleus and DBSCAN clusters. **g** Magnified views centered on Pol II clusters of different sizes. Left: Clustered Pol II localizations with radius of gyration (black dashed circle) and outline considered for counting chromatin nanodomains (cyan dashed circle). Middle: H3K27ac localizations and DBSCAN clusters with centroid. Right: H3K27ac cluster centroids within a distance of <100nm of the Pol II cluster radius of gyration were counted. **h** Schematic illustration of the counting procedure. **i** Number of H3K27ac nanodomain centroids as a function of Pol II cluster radius (magenta) and values obtained at randomized control coordinates (gray, average of 10 randomizations for each cluster). N = 26,902 Pol II clusters. For statistical analysis we binned by Pol II cluster size and compare the number of **k** H3K27ac, **l** H3K4me3 (n = 10,858), and **i** H3K4me1(n = 20,740) nanodomains at Pol II clusters to randomized controls (shaded box plots). Larger Pol II clusters associate with significantly more chromatin nanodomains than randomly expected. One-tailed Wilcoxon rank-sum test. Scale bar: **a,f** 2μm; **a** magnified views 200nm.

We found chromatin nanodomains of similar size for all histone marks in our experiments (**Fig. 5c, Suppl. Fig. 5i**). The fraction of localizations assigned to clusters varies more broadly (**Fig. 5d**). On average, 41% of H3K27ac localizations are assigned to clusters, but 58% of H3K4me3 localizations are clustered. This is consistent with the fact that H3K27ac is typically more pervasive throughout the genome while H3K4me3 is concentrated in sharp peaks at transcription start sites in ChIP-seq profiles ^20^. The inactive H3K27me3 mark clusters to the highest degree (80%) and predominantly in dense regions at the nuclear lamina (**Suppl. Fig. 3d**). Pair-correlation analysis of global localization distribution confirms the DBSCAN-based results (**Fig. 5e**). Notably, chromatin nanodomains are smaller than Pol II clusters (**Suppl. Fig. 5i**) despite the fact that active chromatin spreads far around Pol II clusters (**Fig. 3**).

### Large Pol II clusters are associated with multiple active chromatin nanodomains

We reasoned that multiple individual chromatin nanodomains should associate with single Pol II clusters to explain the wide spread of active chromatin around clusters. To quantify spatial association, we counted the number of active chromatin nanodomains (H3K27ac) located at Pol II clusters (**Fig. 5f**). Given the spread of active chromatin enrichment beyond Pol II cluster boundaries, we considered chromatin nanodomain centroids located in a volume extending 100nm beyond the radius of gyration of each individual cluster (**Fig. 5g,h**). Indeed, multiple active chromatin nanodomains associated with individual Pol II clusters. The number of nanodomains increased with the Pol II cluster radius (**Fig. 5i**). As a control, we randomized the Pol II cluster centroid coordinates in the same nucleus and repeated the analysis. The number of experimentally observed nanodomains diverged from the randomized control for Pol II clusters >100nm. Some large Pol II clusters associated with 10 or more distinct H3K27ac nanodomains.

To facilitate statistical comparison, we again grouped Pol II clusters by size. The average number of distinct H3K27ac nanodomains for Pol II clusters >200nm was 5.5, significantly more than expected from a random distribution (3.0) (**Fig. 5j**). The statistical significance in the enrichment over a random distribution vanished only for small Pol II clusters (50-100nm). H3K4me3 (**Fig. 5k**) and H3K4me1 (**Fig. 5l**) followed the same trend. The number of H3K4me3 nanodomains exceeded that of H3K27ac and H3K4me1 (also see **Suppl. Fig. 5**).

The fact that small Pol II clusters associate with no more active chromatin nanodomains than randomly expected may reflect the fact that at the molecular level, Pol II and nucleosomal epigenetic marks cannot be present simultaneously on the same stretch of chromatin ^40^. Single Pol II and nucleosomes both occupy 50-200bp of double-stranded DNA and are mutually exclusive on those scales. 5-10 polymerases loaded onto the same transcribed element, e.g. in a transcriptional burst, would hence readily result in >= 1kb of nucleosome-depleted chromatin.

### Multiple chromatin elements are co-transcribed in large Pol II clusters

The observation that multiple active chromatin nanodomains associate with single Pol II clusters prompted us to conduct the same analysis for the elongation-specific Ser2p-Pol II phosphorylation mark. Generally, Ser2p signal – as well as Ser5p signal – is distributed throughout the volume of large Pol II clusters (**Suppl. Fig. 6**). We do not find evidence for peripheral localization of elongating marks ^16,23^. While Ser5p initiating Pol II is present throughout the volume of general Pol II clusters (**Fig. 6a**), Ser2p forms distinct foci of smaller size (**Fig. 6b**) in agreement with the analyses presented in **Fig. 1**.

**Fig. 6:**
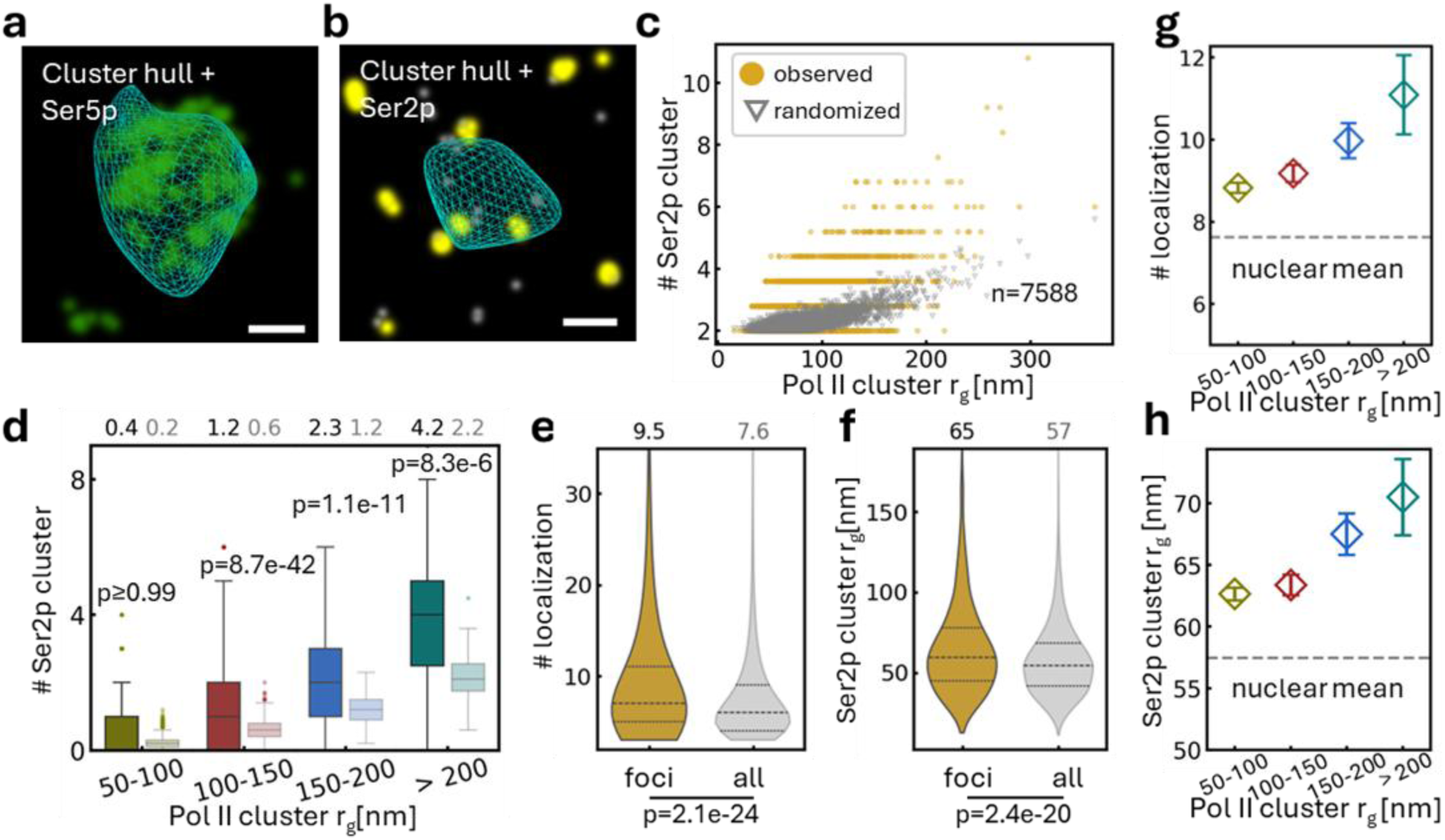
Large Pol II clusters are transcriptional hubs that enhance burst size. Spatial distribution of phosphorylation marks at large Pol II clusters (cyan wireframe). **a** Ser5p initiating Pol II (green) is distributed throughout the cluster volume. **b** Ser2p elongating Pol II (gold) forms distinct foci that may represent transcriptional bursts. Several separated foci are found in single large Pol II clusters. **c** We counted the number of Ser2p foci as a function of Pol II cluster size (gold) and compared to a randomized control (gray). Pol II clusters >100nm radius harbor more Ser2p foci than randomly expected. **d** For statistical comparison, we grouped data by Pol II cluster size. On average, the largest Pol II clusters associate with 4.2 Ser2p foci. **e** The number of super-resolution localizations in Ser2p foci (n=1645) at Pol II clusters of all sizes is significantly larger than the same metric for all Ser2p foci identified throughout the nucleus (n=10,864). **f** Similarly, Ser2p foci at Pol II clusters have a larger radius of gyration. **g,h** Both average number of localizations and radius of gyration of Ser2p foci increase with Pol II cluster size, suggesting that genes at large Pol II clusters produce larger transcriptional bursts. Dashed lines indicate the mean values for Ser2p foci across the entire nucleus. One-tailed Wilcoxon rank-sum test. Scale bar: **a** 200nm.

We counted the number of Ser2p foci associated with each general Pol II cluster with the same approach as described above for chromatin nanodomains. Multiple clearly distinct Ser2p foci associate with single large Pol II foci. The number of Ser2p foci increases with Pol II cluster size (**Fig. 6c**). The largest Pol II clusters typically harbor >5 distinct Ser2p foci. The difference to randomly expected numbers is again highly significant for Pol II clusters >100nm (**Fig. 6d**) and comparable to the number of active chromatin nanodomains at Pol II clusters of the same size. Together, the analyses of active chromatin nanodomains and elongating Pol II foci suggest that Pol II clusters reflect hubs of co-transcribed active chromatin elements.

### Association with Pol II clusters increases transcriptional burst size

Transcriptional bursts of the stem cell transcription factor Sox2 increase in proximity to transcription condensates, demonstrating a functional impact of condensates on gene regulation ^15^. We hypothesized that this effect would be reflected in Ser2p data if it transfers to other genes in contact with Pol II clusters. Assuming that Ser2p foci represent trains of elongating polymerases, i.e. transcriptional bursts, we determined whether transcriptional burst size changes in the vicinity of Pol II clusters. To assess a possible impact of Pol II clusters on transcriptional bursts, we compared Ser2p foci properties at Pol II clusters to properties of Ser2p foci everywhere in the nucleus. We evaluated the number of localizations (**Fig. 6e**) and radius of gyration (**Fig. 6f**). Ser2p foci and hence transcriptional bursts at Pol II clusters are significantly larger than the nuclear average by both metrics. Moreover, Ser2p foci size grows with Pol II cluster size (**Fig. 6g,h**). Ser2p foci are sparse and clearly spatially distinct (**Fig. 6b**). Hence, the size increase is not an artifact of overlapping Ser2p foci. This finding is direct evidence for increased functional output through cooperative action as predicted by the transcription condensate model ^13^ .

### A chromatin hub model for Pol II clustering and cooperative activity

We propose that Pol II clusters reflect active chromatin micro-environments, specifically hubs of multiple transcribed elements that permit a large number of polymerases to be loaded onto chromatin. Accordingly, we observe multiple distinct active chromatin nanodomains associated with individual Pol II clusters. Ser5p cluster properties are globally indistinguishable from those of general Pol II clusters, and Ser5p signal fills the volume of general Pol II clusters as previously reported ^14^. In contrast, multiple distinct Ser2p foci associate with single general Pol II clusters. This is reasonable since genes are often transcribed in bursts while the transition of Pol II from chromatin loading through initiation and promoter proximal pausing into productive elongation is very inefficient ^35^. In our interpretation, the larger accumulation of Ser5p signal in general Pol II clusters mirrors the abundance of H3K4me3 chromatin found at transcription start sites of multiple promoters. The presence of multiple distinct Ser2p foci reflects co-transcription of chromatin elements in the hub structure ^41^, which appears to make the transition into elongation more efficient and increases transcriptional burst size as measured by Ser2p foci properties. It is well documented that gene promoters can also act as enhancers of transcription for other genes ^42^. Notably, more stereotypical enhancer marks (H3K4me1) associate peripherally with the promoter hubs. This raises the question of what orchestrates the spatial organization of chromatin into such structures.

### Chromatin hubs at condensates are resistant to transcription inhibition

The coalescence of active chromatin elements decorated by BRD4 has been proposed as a mechanism to link distal elements via condensate formation ^19,21^. Knowing that the largest Pol II clusters are rich in BRD4 ^7,12^, we incubated cells with the small molecule bromodomain inhibitor JQ1 (1 μM, 90 min). JQ1 prevents BRD4 binding to acetylated chromatin ^43^. We quantified chromatin mark enrichment at large Pol II clusters that are detectable by widefield fluorescence microscopy (**Fig. 7a**). We also included a label for the Mediator subunit MED1, a second prominent condensate component. Pol II and MED1 colocalize in transcription condensates. We labeled both proteins in addition to histone marks to avoid potential issues with protein redistribution under drug treatment. To this end, we used a homozygous Snap-Pol II/Halo-MED1 cell line generated from the parental v6.5 line used above ^14^. Endogenous Pol II and MED1 were labeled with Snap-JFX 650 and Halo-TMR ligand, respectively, and tertiary marks were stained with an AF488-labeled antibody system. Under JQ1 treatment, we indeed observed a loss of H3K27ac enrichment at Pol II condensates (**Fig. 7b**). But surprisingly JQ1 treatment did not result in the loss of other chromatin marks at condensates. We observed a minor reduction in H2B enrichment but H3K4me1 and H3K4me3 levels were virtually unchanged. H3K27me3 levels around condensates dropped slightly, which may indicate that H3K27ac chromatin that is lost from the condensate core infiltrates this surrounding region. Our observations agree with the proposed ability of BRD4 to tether acetylated chromatin to condensates but also indicate that inhibition of BRD4 binding does not otherwise perturb the chromatin structure at condensates. To ensure that the observed effects reflect local changes at Pol II condensates and not absolute changes we quantified absolute nuclear intensities in all experiments and compared to the untreated condition (**Suppl. Fig. 7**).

**Fig. 7:**
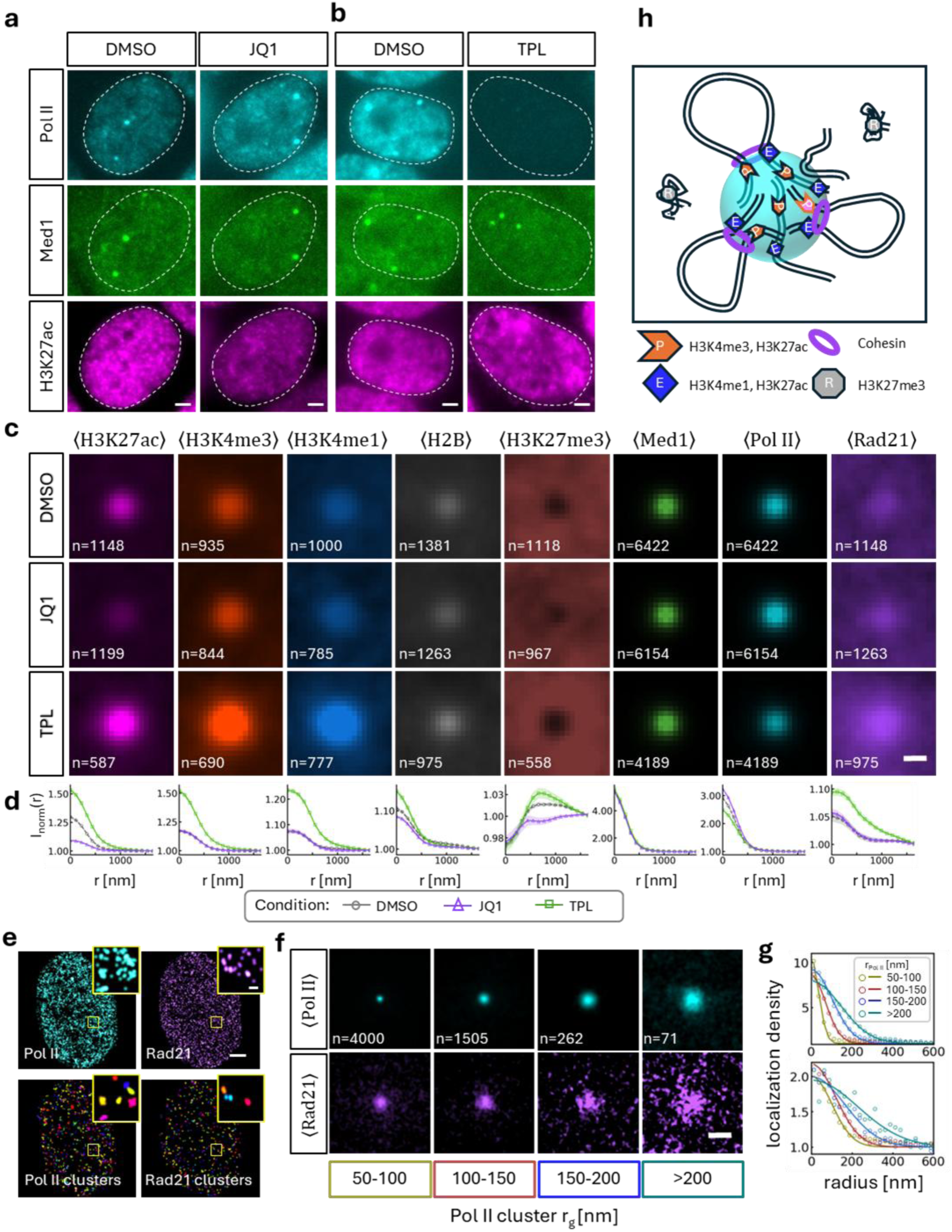
Chromatin structures contract in the absence of Pol II. **a** Conventional resolution three-color fluorescence microscopy images of the condensate constituents Pol II and Med1 as well as the active chromatin mark H3K27ac under JQ1 and **b** triptolide (TPL) treatment. Prolonged TPL exposure leads to Pol II degradation while Med1 is retained in condensates. **c** Assessment of chromatin structural changes at condensates upon JQ1 and TPL incubation by accumulation of signal at condensates and **d** relative radial intensity at condensates normalized to average signal in the nucleus. JQ1 (purple triangles) reduces H3K27ac enrichment while other marks are mostly unchanged. TPL (green squares) increases relative enrichment of all active chromatin marks and H2B while Med1 enrichment is unchanged. Absolute abundance and enrichment of Pol II is reduced by 80% (Fig. 1b), but relative enrichment is only slightly reduced. **e** Super-resolution images of Pol II and the cohesin subunit Rad21 as well as cluster identification by DBSCAN. **f** Rad21 is enriched at Pol II clusters of all sizes to similar degrees. **g** At larger Pol II clusters, the spread of Rad21 but not its concentration increases.

Other studies suggest that the Pol II C-terminal domain itself has self-affinity that allows it to form clusters or condensates in the nucleus through homotypic interactions ^29,44,45^, though our own experiments demonstrate that this is not the cause of Pol II accumulation in condensates ^14^. To remove Pol II from promoter chromatin, we exposed cells to the initiation inhibitor triptolide for extended periods of time. Prolonged triptolide exposure leads to proteasome-dependent Pol II degradation. 90 min incubation with 10 μM triptolide resulted in loss of 80% of Pol II nucleus-wide (**Fig. 7b, Suppl. Fig. 7**). We observed reduction of absolute Pol II levels in the nucleus, but relative enrichment in condensates remained unchanged (**Fig. 7c**). To detect condensates reliably under conditions with reduced Pol II levels, we relied on Mediator signal (MED1). The Mediator complex colocalizes with Pol II in condensates ^7^ . Loss of Pol II did not impact the MED1 enrichment in condensates such that transcription condensates remained detectable in this channel despite Pol II degradation (**Fig. 7b**) as previously reported ^14^. Under initiation inhibition and Pol II degradation, the relative enrichment of all active chromatin marks in condensates at least doubled (**Fig. 7c,d**). H2B enrichment also increased markedly.

We conclude that neither chromatin binding of Pol II nor active transcription are the driver of chromatin hub formation. On the contrary, depletion of Pol II allows the underlying chromatin structure to compact. These findings are reflected in global changes of chromatin association with condensates detected by sequencing [*Bogdanovi*ć *et al.: Large RNA polymerase II condensates are promoter-centric assemblies.* Reference to be inserted].

Based on these observations, we hypothesized that chromatin architectural factors may drive the formation of chromatin hubs. Prior reports indicated a loss of transcription condensates upon CTCF degradation ^46^. CTCF is considered a roadblock for cohesin-mediated DNA loop extrusion. To determine whether cohesin might be responsible for forming active chromatin topologies that recruit transcription condensates, we stained for the cohesin subunit Rad21 (**Fig. 7e**). Indeed, we observed relative enrichment of Rad21 at condensates that is unchanged under JQ1 but increases under triptolide treatment and loss of Pol II (**Fig. 7c,d**). It has been shown that Pol II transcription presents a roadblock to cohesin loop extrusion ^47,48^. Enrichment of cohesin at condensates is consistent with a model in which DNA loop extrusion acts to create the active chromatin hubs underlying Pol II condensates but is restrained by the presence of Pol II ^49^. We followed up on these initial observations with super-resolution microscopy to determine if Rad21 enrichment persists at smaller Pol II clusters (**Fig. 7e**). These experiments confirm that Rad21 is enriched at Pol clusters of all sizes to a similar degree (**Fig. 7f**). As reported for active chromatin marks (**Fig. 3**), Rad21 enrichment extends beyond Pol II cluster boundaries and the spread of Rad21 enrichment increases with Pol II cluster size (**Fig. 7g**). This observation is consistent with a model in which chromatin hubs formed by loop extrusion serve as the substrate for spatial accumulation of Pol II (**Fig. 7h**).

## DISCUSSION

Spatial and temporal clustering of transcription has been studied for a long time. We find that spatial clusters of Pol II are rich in Serine 5-phosphorylated, initiating Pol II. Elongating Ser2p Pol II clusters are relatively more sparse and smaller. This mirrors earlier studies that showed the transition from promoter loading through initiation and pausing into productive elongation to be a highly inefficient process with only 1% success rate ^35^. In an accompanying submission we show that condensate increase the efficiency of this process and are rich in highly expressed genes [*Bogdanovi*ć *et al.: Large RNA polymerase II condensates are promoter-centric assemblies.* Reference to be inserted]. We and others have shown that only a minority fraction of Pol II in a nucleus is in an unbound state at a given time ^14,35^.

In agreement with a model in which most Pol II is engaged with chromatin, we find that Pol II clusters form at regions of high active chromatin density. Vice versa, active chromatin enrichment spreads to a characteristic size of a few 100nm beyond Pol II cluster boundaries. The size of these active chromatin domains is reminiscent of chromatin regions identified in nuclear cryosections by electron microscopy ^37^. It also corresponds to the approximate range within which regulatory contacts can be established based on high resolution chromatin tracking ^50^, and the range over which transcription condensates have been shown to increase Sox2 transcriptional burst size ^15^. The active chromatin enrichment domains grow in size, but not in density for larger Pol II clusters. The fact that active chromatin enrichment extends well beyond the size of Pol II clusters indicates that these chromatin organizational structures exist independent of Pol II clusters. Our results suggest that Pol II clusters represent protein recruitment to these chromatin scaffolds. Notably, H2B nucleosomal chromatin density is also locally enriched at the coordinates of Pol II clusters of all sizes, i.e. endogenous transcription condensates do not displace chromatin.

More specifically, we observed that H3K4me3 which is found at transcription start sites enriched more within Pol II clusters than enhancer marks. H3K4me1 localization density is the highest precisely at the surface of large Pol II clusters. With super-resolution methods we were able to count multiple spatially distinct active chromatin clusters overlapping with single Pol II clusters. Thus, Pol II clusters form at active chromatin hubs in which multiple regulatory elements, specifically H3K4me3-decorated promoters, colocalize. This explains why general Pol II clusters have almost the same properties as Ser5p clusters, and why with single particle tracking we observe a high abundance of chromatin-bound Pol II in large Pol II clusters ^14^. The largest Pol II clusters, transcription condensates, are thought to associate with super-enhancer controlled gene loci ^13^. A recent study showed that super-enhancers are 3D spatial hubs of colocalizing genes ^26^. The presence of multiple active chromatin elements at large Pol II clusters agrees with this model, although our super-resolution data suggests that the assemblies are promoter-centric as predicted from the function of Pol II as a roadblock to cohesin loop extrusion ^49^. Prior studies had shown association of specific, often super enhancer-controlled gene loci with transcription condensates but lacked the spatial and DNA sequence resolution to distinguish which regulatory elements associated with condensates.

For two specific genes, Esrrb ^7^ and Sox2 ^15^, we were able to show that gene loci associate only transiently with stable Pol II condensates. In the case of Sox2, neither the promoter nor the super-enhancer locus control region permanently overlapped with the nearest condensate ^15^. If condensates indeed harbor multiple colocalized promoters, one can rationalize these observations. Any specific one of the promoters may associate with the condensate chromatin hub only transiently, but the structure can persist without all of its members present. This description converges with the original model of transcription factories that act on genes passing through them.

The presence of multiple Ser2p foci at large Pol II clusters indicates that clustered genes are co-transcribed at these sites. Using Ser2p foci properties as a proxy for transcriptional bursts, we find that burst size at condensates is significantly increased. We had previously characterized this effect for a specific gene, Sox2 ^15^. The data presented here provides for the first time evidence that this effect is not linked to select genes but represents a universal mechanism. Accordingly, a new assay to extract condensates from cells and identify chromatin elements associated with them shows association of many genes with these nuclear bodies [*Bogdanovi*ć *et al.: Large RNA polymerase II condensates are promoter-centric assemblies.* Reference to be inserted]. Purified condensates are rich specifically in promoter and proximal enhancer chromatin of many highly expressed genes.

The cooperative effect of enhanced transcriptional bursts increases with Pol II cluster size and hence with the number of colocalized transcribed elements, possibly because active promoters themselves can act as enhancers for other promoters ^51^. Such cooperative effects are expected from the transcription condensate or factory models.

Assembly of the chromatin structures may be driven by cohesin-mediated loop extrusion. Upon transcription inhibition and Pol II degradation, chromatin and Rad21 signatures at the site of condensates compact. Others had previously shown that CTCF deletion leads to disassembly of condensates ^46^. Perturbation of condensates forming proteins (BRD4) with a suspected role in chromatin organization led to a reduction of H3K27ac, but not other marks at condensates.

## RESOURCE AVAILABILITY

### Lead contact for reagent and resource sharing

Further information and requests for resources and reagents should be directed to and will be fulfilled by the lead contact, Dr. Jan-Hendrik Spille.

### Materials availability

Reagents generated in this study are available from the lead contact upon request.

### Data and code availability

[Analysis code will be made publicly available through the Spille Lab Github repository.] Any additional information required to reanalyze the data reported in this paper is available from the lead contact upon request.

## ACKNOWLEDGEMENTS

We thank members of the Ruthenburg Lab (UChicago) and Dr. Primal de Lanerolle for helpful discussions. We gratefully acknowledge support from the Research Corporation for Scientific Advancement (CMC 28407), NIH (R35GM150560), NSF (MCB-2306187), and the University of Illinois Chicago Startup Fund. This research was supported in part by grants from the NSF (DMS-2235451) and Simons Foundation (MP-TMPS-00005320) to the NSF-Simons National Institute for Theory and Mathematics in Biology (NITMB).

## AUTHOR CONTRIBUTIONS

Conceptualization: G.P., J.G.-S., and J.H.S. Methodology: G.P., J.G.-S., A.B., F.M., and J.H.S.

Investigation: G.P., J.G.-S., A.B. Formal analysis: G.P., J.G.-S. Software: G.P., J.G.-S.

Visualization: G.P., J.G.-S., A.B. and J.H.S. Funding acquisition: J.H.S. Project

administration: J.H.S. Supervision: J.H.S. Writing – original draft: G.P., J.G.-S., and J.H.S.

Writing – review & editing: G.P., J.G.-S., and J.H.S.

## DECALARATION OF INTERESTS

The authors declare no competing interests.

## DECALARATION OF GENERATIVE AI AND AI-ASSISTED TECHNOLOGIES

The authors declare no competing interests.

## SUPPLEMENTARY INFORMATION

Supplementary Figures 1-7

Supplementary Tables 1-2

## METHODS

### Cell culture and immunostaining

Mouse embryonic stem cells (v6.5) were maintained under 2i culture conditions using the ESGRO 2i medium kit (SF016-200, Sigma-Aldrich) supplemented with 15% fetal bovine serum (EmbryoMax ES Cell Qualified FBS). Cells were cultured in 24-well plates precoated with 0.2% gelatin for 30 min at 37°C in a humidified incubator containing 5% CO₂.

Cells were seeded onto either 35 mm glass-bottom dishes (Celvis D35-20-1.5H) or 8-well chambered coverglass slides (Celvis C8-1.5H-N). The imaging dishes were freshly coated with poly-L-ornithine (PLO; Sigma P4957, diluted 1:1 in water) overnight, followed by laminin coating (Sigma L2020, 0.5 µg/mL) for 4 h prior to cell plating.

The next day, samples were fixed either with 3.2% paraformaldehyde (PFA) for 10 min at room temperature, followed by three washes with room-temperature PBS, or with 100% ice-cold methanol for 7 min at −20°C, followed by three washes with cold PBS (4°C) for 5 min each. Cells were permeabilized with 0.1% Triton X-100 in PBS for 10 min and subsequently washed three times with PBS in methanol fixed cases. For PFA-fixed samples, 0.5% Triton X-100 was used. Samples were then blocked for 1 h at room temperature in 4% BSA prepared in PBS. Primary antibodies diluted in 4% BSA were incubated with samples overnight at 4°C in a humidified chamber. The following day, cells were washed three times with PBS (5 min each) and incubated with the appropriate secondary antibodies for 1 h at room temperature. Samples were subsequently washed three additional times with PBS before imaging. Antibody information and concentrations are listed in **Supplementary Table 1** and combination used for different experiments in **Supplementary Table 2**.

### Drug treatments (TPL and JQ1)

Drug treatments were conducted on a homozygous Snap-Pol II/Halo-Med1 cell line created by CRISPR insertion of fluorescent tags in the parental v6.5 line as previously described ^14^. This allowed for direct labeling of endogenous protein. Cells were incubated with JFX650 Snap (50nM) and Halo TMR (200nM) for 20 min and washed 2X before incubation with inhibitors. Cells were fixed with ice-cold methanol for 6 minutes, followed by 3 times cold PBS wash and immuno-staining as described.

We prepared a 1000x stock concentration of triptolide (TPL, MedChemExpress, HY-32735) and (+)-JQ-1 (MedChemExpress, HY-13030), respectively in DMSO. Stocks were aliquoted to 10 µl, and stored at -80 °C until the day of the experiment. Cells were incubated with TPL/JQ1 at a final concentration of 10 µM TPL/1 µM JQ1 for 90 min after staining with Halo- /Snap-ligands. As a control, separate dishes were incubated with an equivalent volume fraction of DMSO.

### Super-Resolution Microscopy

Super-resolution imaging buffer was freshly prepared before imaging as previously described ^52^. Briefly, 30 µL of OxyFluor (Oxyrase Inc., Mansfield, OH, USA) was mixed with 680 µL of ice-cold PBS (4°C). Subsequently, 50 µL of sodium DL-lactate (Sigma #L1375) was added and vortexed, followed by the addition of 50 µL of 1 M MEA (Sigma #30070) to achieve a final concentration of 50 mM MEA. Finally, 40 µL of NaOH was added to adjust the pH to approximately 8.0. The imaging buffer was filtered through a 0.2 µm syringe filter and maintained on ice throughout preparation.

Bi-plane ^32^ super-resolution imaging was performed on a Bruker Vutara 352 microscope. Bruker SRX software was used for acquisition of 3d dSTORM localizations. Sequential imaging was performed by first exciting Alexa Fluor 647 with a 640 nm laser (5.3 kW/cm²), followed by Alexa Fluor 488 excitation using a 488 nm laser (5.0 kW/cm²). For each fluorescence channel, 50,000 image frames were collected with an exposure time of 20 ms per frame.

For drift correction and chromatic alignment, 100 nm gold fiducials (TED PELLA, Cat#15711-20) were used. Fiducial stock solution was diluted 1:8 in PBS, briefly vortexed, sonicated for at least 10 min, and vortexed again immediately prior to application. The diluted fiducial solution was added to the sample and incubated for 10 min, after which it was aspirated, and the samples were washed twice with PBS before the addition of super-resolution imaging buffer.

Localizations were filtered in the SRX software to remove low-confidence detections. Drift and chromatic aberration corrections were applied using built-in functions. Individual nuclei were manually segmented, and localization datasets were exported for downstream analysis in Python.

### Cluster identification

Clusters were identified using the DBSCAN function implemented in scikit-learn^53^. DBSCAN parameters were selected empirically based on visual inspection of clustering behavior and were kept consistent across channels. General Pol II and Ser5P-Pol II localizations were analyzed using eps=100 nm and min_samples= 5, Ser2P-Pol II localizations using eps=110 nm and min_samples= 4, and chromatin marks using eps=100 nm and min_samples= 4.

In order to generate a 3-dimensional nuclear hull, localizations from both imaging channels were combined, and an alpha surface was computed using the alpha_wrap_3 function from Python bindings of the CGAL^54^ with an alpha radius of 500nm and an offset distance of 20nm.

To reduce variability arising from differences in localization density across datasets, localizations were randomly subsampled within each nucleus prior to clustering analysis. Final target densities were set to 139 localizations/µm³ for general Pol II and Ser5P-Pol II, 83 localizations/µm³ for Ser2P-Pol II, 73 localizations/µm³ for Rad21, and 100 localizations/µm³ for all chromatin marks. Nuclei with less than the indicated density in any channels were removed from the analysis.

### Predictivity measurement

Chromatin mark density was estimated for each nucleus using kernel density estimation implemented with the KernelDensity function from scikit-learn using a bandwidth of 100 nm. Density distributions for each nucleus were normalized by the mean density of that nucleus. Mark density values were then evaluated at each Pol II cluster centroids, and 10,000 random points were sampled within each nuclear volume to establish a random reference. Pol II clusters were subsequently grouped into 20 bins based on normalized chromatin density, and the observed number of clusters in each bin was compared with the expected number based on the random sample.

### Chromatin mark spatial enrichment analysis

For radial aggregation analysis, 3d localizations were aligned with reference to Pol II cluster centroids in both the Pol II and the chromatin mark channel. Localizations within a z-extent equal to the Pol II radius of gyration were included. Radial localization densities were computed in 2D from a random sample of at most 4000 clusters. If the cluster size class had fewer than 4000 clusters, all were considered. The values are divided by the mean background localization density, measured in an annulus region of 500 nm – 1000 nm. For surface enrichment analysis in 3D, a surface hull enclosing each Pol II cluster was generated using the Python bindings of the CGAL alpha_wrap_3 function with an alpha radius of 200 nm and an offset of 20 nm. Localization counts were measured at distances up to +600 nm from the cluster surface in 20 nm intervals. The innermost radius considered was adjusted depending on the Pol II cluster radius of gyration (-100nm for clusters >200nm, -80 nm for 150nm-200nm, -60 nm for 100-150 nm, and -40 for Pol II clusters of size 50 nm-100nm). Resulting densities were smoothed using a moving average with a window size of 3. To estimate the random expectation, each Pol II cluster hull was randomly repositioned 10 times within the same nuclear volume, and the mean localization count from these random placements was calculated. Enrichment scores were computed as the ratio of experimentally observed to randomly expected localization counts. Confidence intervals were estimated by bootstrapping (10 times resampling the clusters with replacement). Enrichment ratio was computed from resampled values. Shaded regions indicate 95% confidence interval values from the bootstrap distribution.

Nanodomains or foci associated with Pol II clusters were identified by counting centroids that fell within a spherical region defined by the 95th percentile cluster radius plus an additional 100 nm buffer centered on each reference cluster.

### Pair and cross-pair correlation analysis

Pairwise distances within or between probe populations were measured in 2D using 2 nm radial bins. Neighbor searches were performed using the cKDTree function from SciPy^55^. Observed pair counts were normalized against randomized point distributions sampled within the same nuclear hull, and correlation functions were calculated as the ratio of observed to expected pair counts.

### Condensate segmentation in 3-color widefield images

We use the Med1-Halo-TMR signal to identify condensates in both DMSO and drug-treated cases. First, each image was normalized by subtracting the mean and dividing by the intensity range (computed as the difference between the 99th percentile and the 1^st^ percentile in order to exclude extreme outliers/bad pixels). Image were filtered using scipy’s Laplacian of Gaussian, and binarized by thresholding. Condensate peaks were identified from connected binarized regions based on area, relative intensity, and eccentricity. The signal from all three color channels for each region of interest was aligned by the condensate centroid. The same thresholding and filtering parameters were used across all experiments. Nucleolar regions were identified using the ‘2D_versatile_fluo’ pretrained model of Stardist2D on the Pol II -Snap-JFX650 channel followed by scikit-image’s hessian filter and removed from the nuclear mask.

## Notes

### Competing Interest Statement

The authors have declared no competing interest.

