## Supplementary Information for "Transcription condensates are promoter hubs that enhance transcriptional bursts"

Supplementary figures 1-7

Supplementary tables 1-2

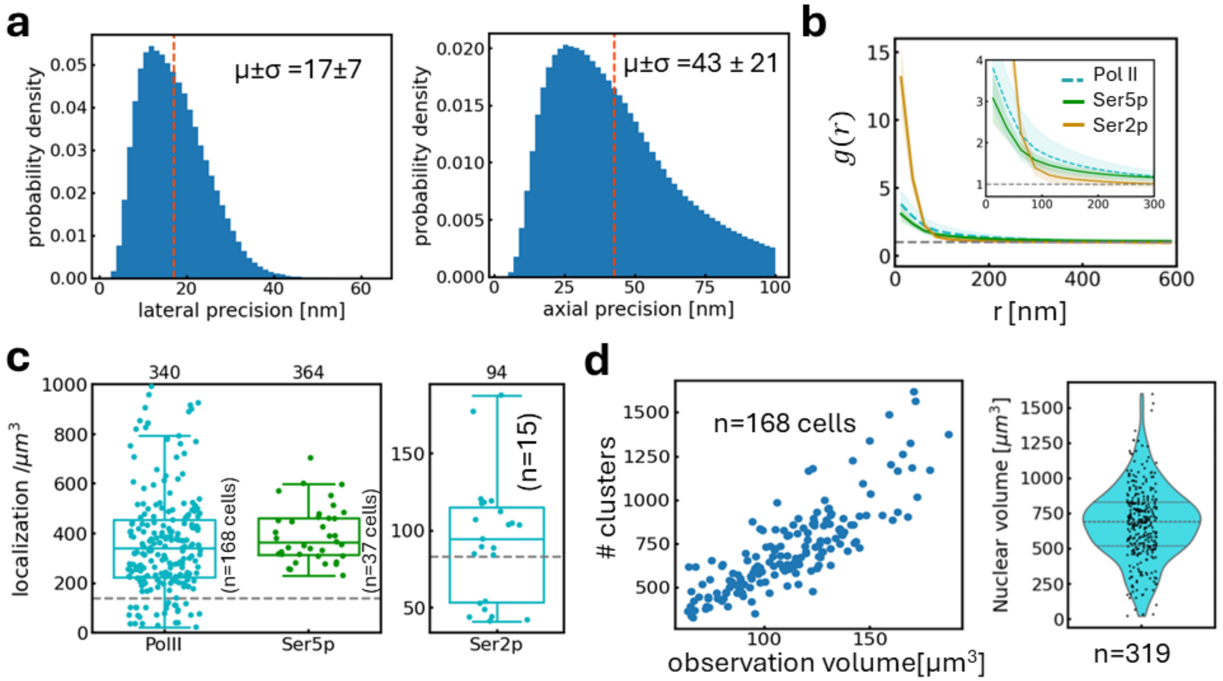

**Suppl. Fig. 1: RNA Pol II cluster properties**.

a) Lateral and axial localization precision distribution in biplane 3D super-resolution microscopy calculated from camera, point spread function, and number of photons using the SRX microscope control software (Bruker).

b) Pair correlation analysis based on all localization without subsampling random thinning. The results do not differ from Fig. 1g determined with random thinning.

c) Localization density of general Pol II and Ser5p (left) and Ser2p (right) for all cells in the dataset. Cells with localization density smaller than the threshold value (dashed line) were not considered for analysis to avoid artifacts arising from localization density differences.

d) The number of Pol II clusters identified by DBSCAN increases linearly with the observation volume per cell nucleus. Nuclear volume distribution of WT v6.5 nuclei based on segmentation of Pol II immunostaining in z-stacks. Mean ± sd: 678 ± 262 $\mu m^3$. Based on segmentation results, nucleoli take up 20% of this volume. Since Pol II is excluded from nucleoli, we disregarded this subvolume for the estimate of total clusters per cell.

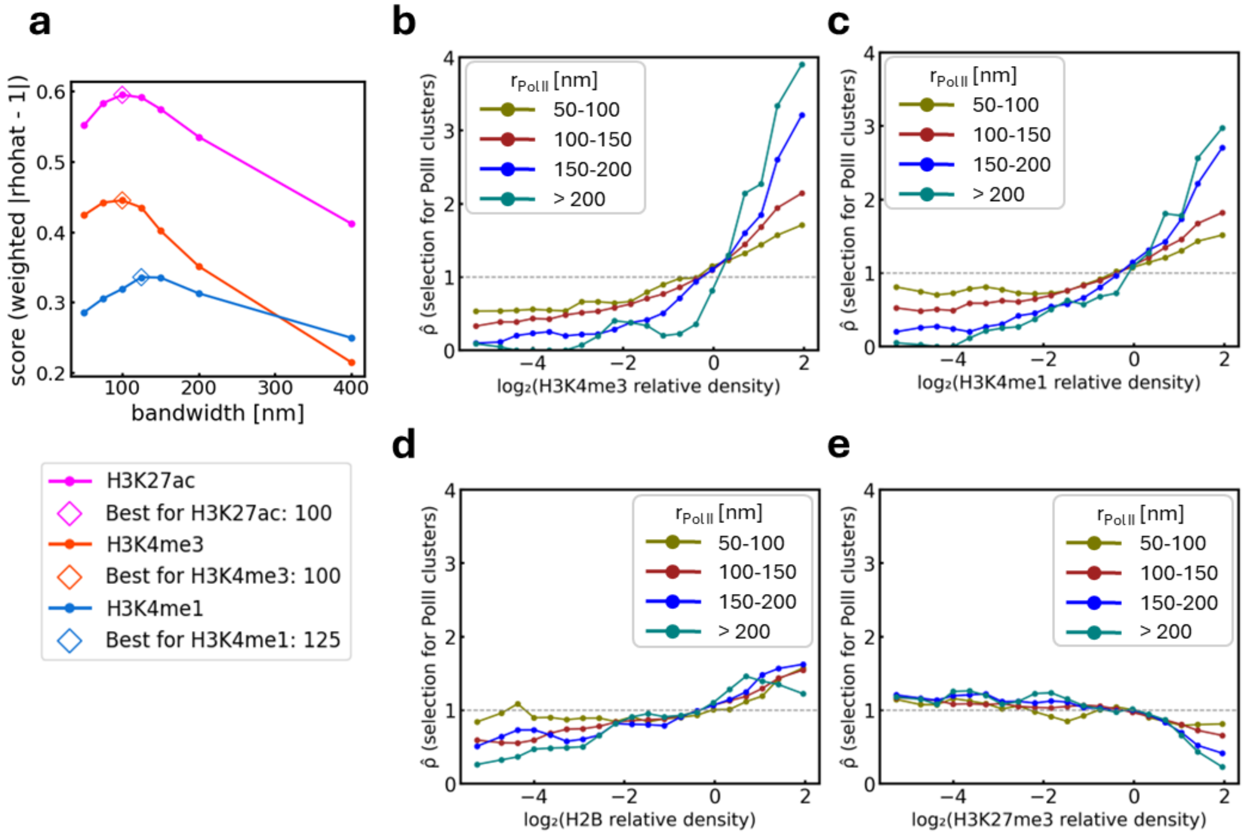

**Suppl. Fig. 2: Predictivity analysis and bandwidth optimization.**

a) Predictivity score as a function of kernel density estimation (KDE) bandwidth. The predictivity score quantifies how strongly the predictivity deviates from random expectation. We computed for each chromatin mark density bin the deviation of $\widehat{\rho}$ from 1 and weighted by the fraction of Pol II clusters within that bin. The weighted deviations are then summed to obtain a global predictivity score of the respective chromatin mark. Higher scores indicate better predictivity of Pol II cluster location. We used this approach to determine that a kernel density of 100nm is optimal. H3K4me1 yielded a slightly larger optimal value, but we analyzed all data with the same kernel density of 100nm.

b) $\widehat{\rho}$ as a function of H3K4me3 density for Pol II clusters of different sizes. Same for c) H3K4me1, d) H2B, and e) H3K27me3 density.

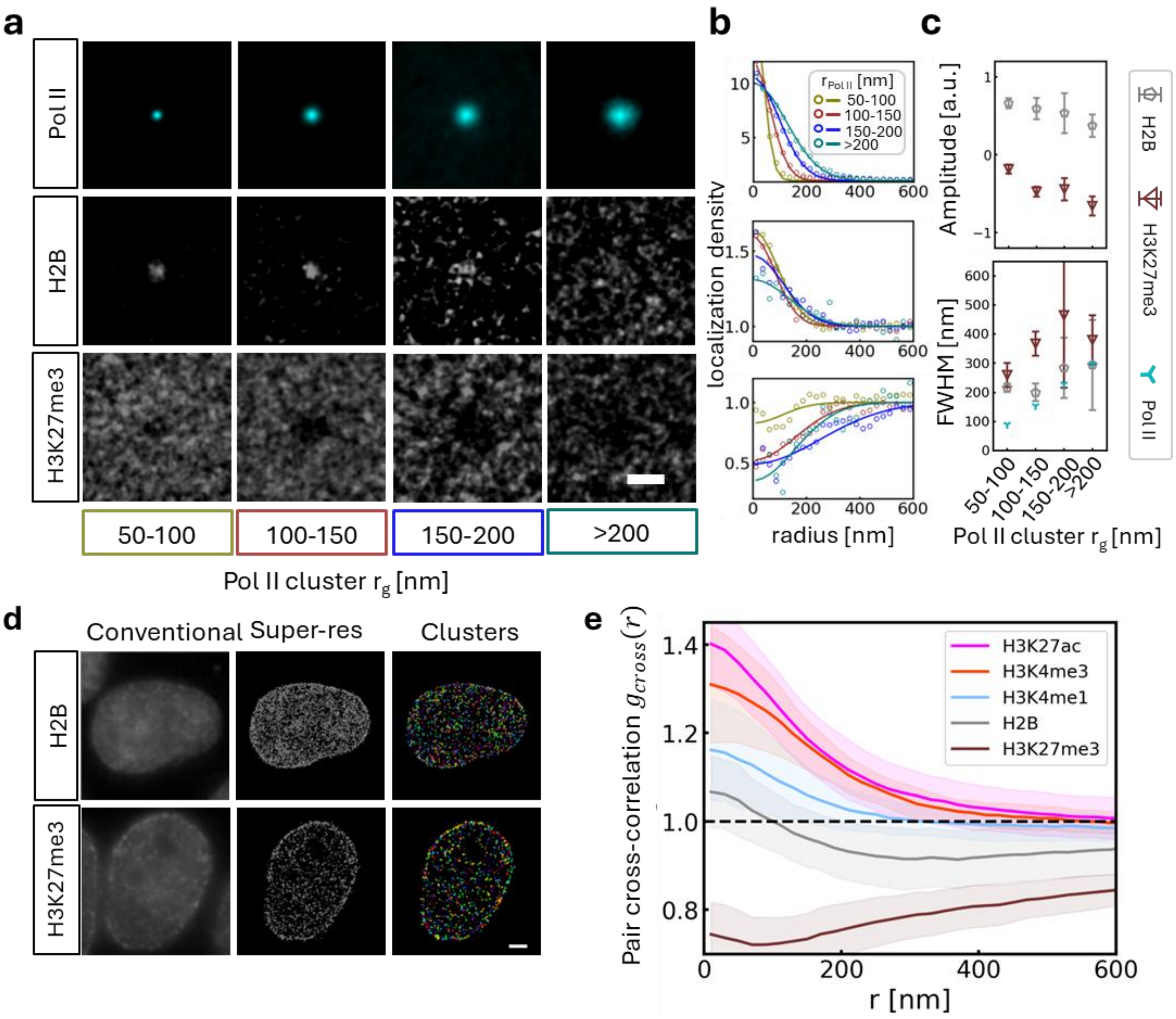

**Suppl. Fig. 3: Chromatin marks at Pol II clusters.**

a) Aggregate localizations at Pol II clusters of different sizes (top) and H2B (middle) or H3K27me3 (bottom) localizations aligned by Pol II cluster centroid. Scale bar 500nm

b) Radial localization density around Pol II cluster centers normalized to the mean density in an annulus of 500nm-1000nm. Circles represent the measured values, lines represent a gaussian fit to Pol II (top), H2B) (middle), and H3K27me3 (bottom) localization density.

c) Fit parameters amplitude and spread (FWHM) obtained from b). Mean and standard deviation from bootstrapping (10 iterations).

d) Conventional (left), and super-resolution (middle) image of H2B and H3K27me3 along with the clusters detected using DBSCAN. Scale bar: 2μm.

e) Pair cross-correlation of histone mark with RNA Pol II localizations.

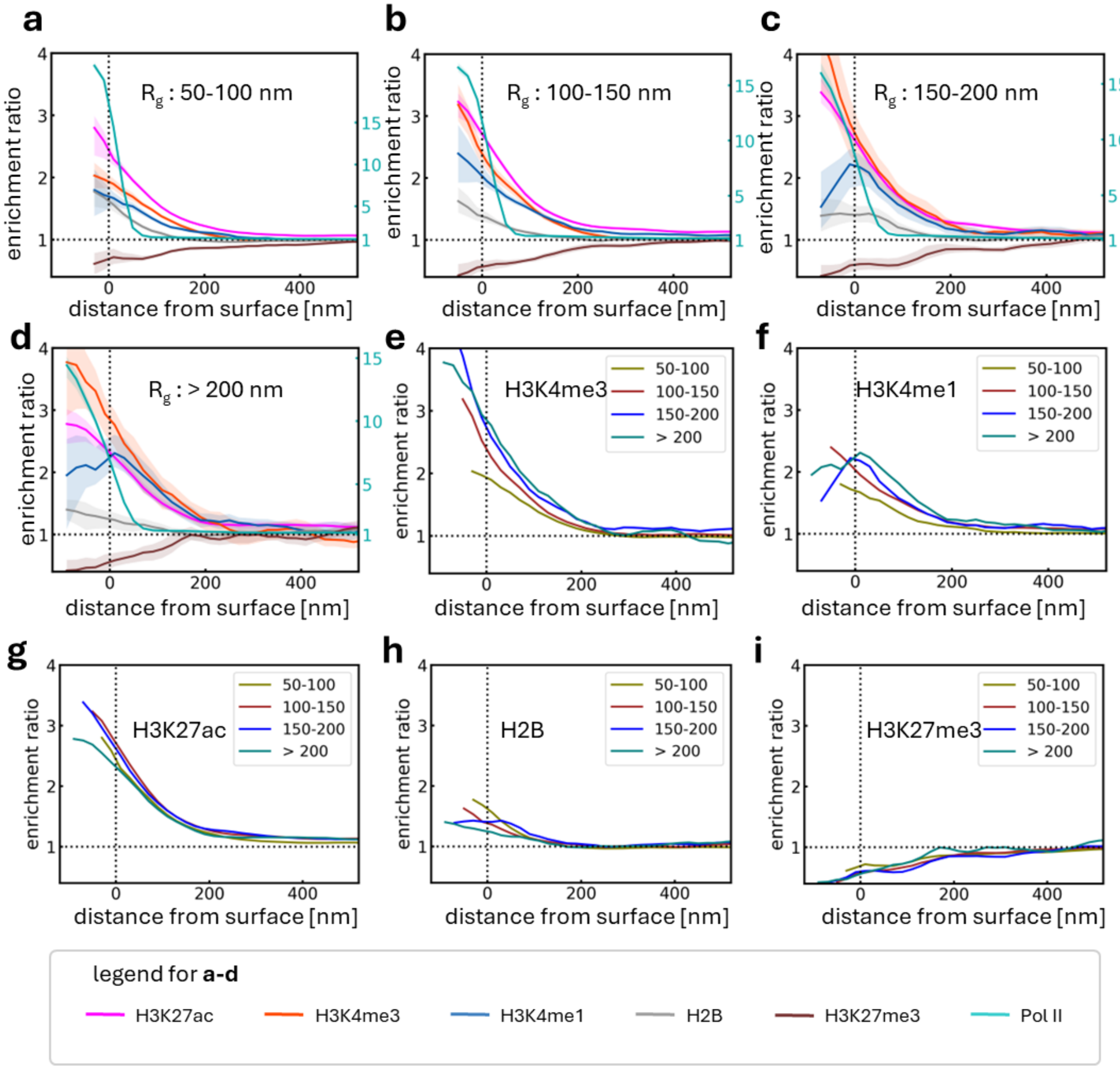

**Suppl. Fig. 4: Cluster size-dependent surface enrichment analysis for different chromatin marks.**

a-d) Enrichment ratio for different chromatin marks as a function of distance from Pol II cluster surface. Negative distance indicates the interior of the cluster. Left axis refers to histone mark enrichment, right axis to Pol II enrichment. The annotated $R_g$ represents the Pol II cluster size range considered for analysis in the respective panels.

e-i) The same enrichment curves sorted by histone mark to compare profiles for Pol II clusters of different size. Legends refer to Pol II cluster radius of gyration bins.

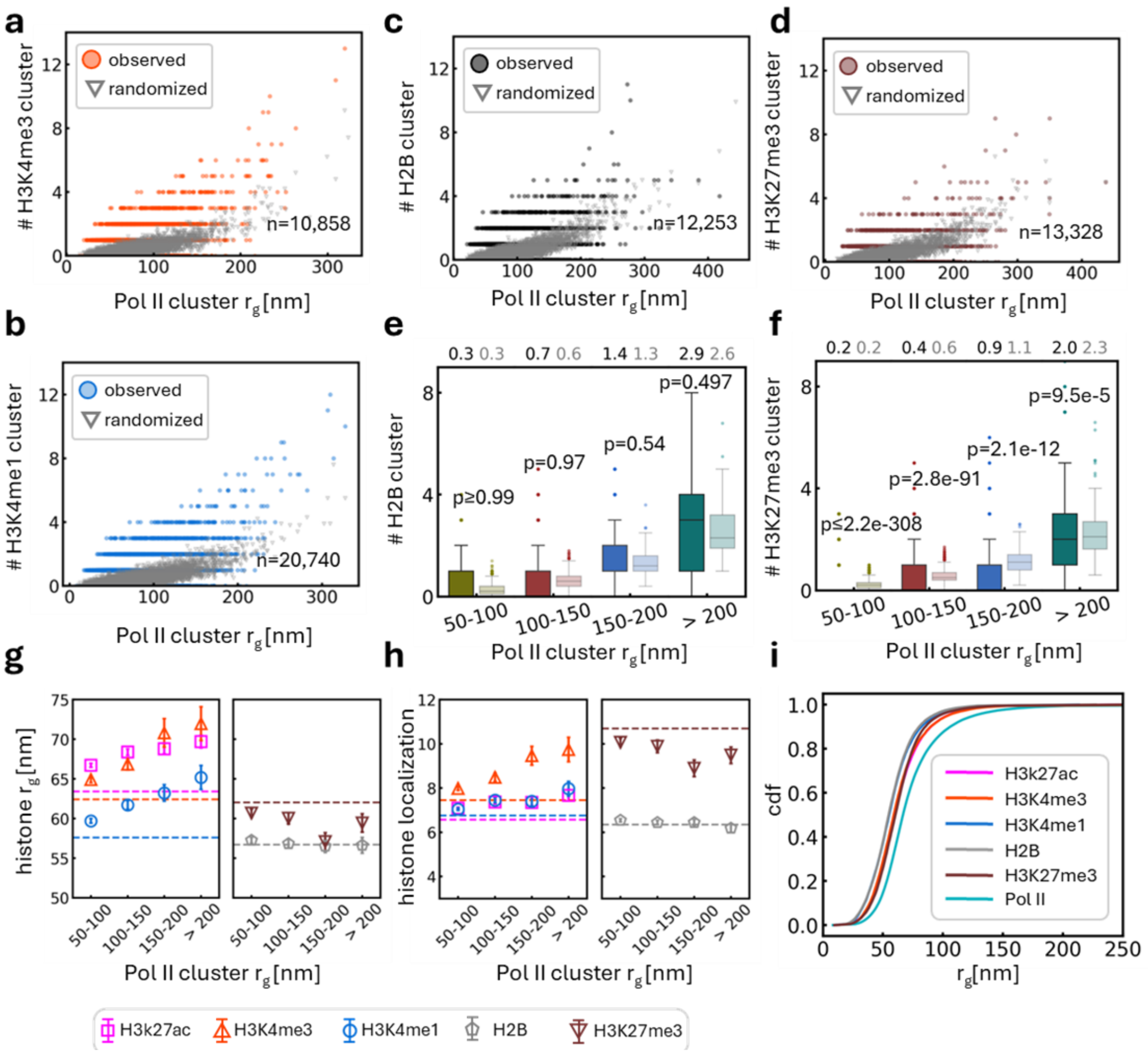

**Suppl. Fig. 5: Chromatin mark cluster properties**

a) Number of H3K4me3 nanodomains vs radius of each Pol II cluster comparing experimental data to results at randomized coordinates (mean of 10 positions per cluster). Same representation for b) H3K4me1, c) H2B, and d) H3K27me3.

e) Mean number of H2B and f) H3K27me3 domains per Pol II clusters in different size bins. One-tailed Wilcoxon rank-sum test.

g) Average nanodomain size plotted against the Pol II cluster radius for active histone marks (left) and controls H2B and H3K27me3 (right). Dashed lines represent the average size of these domains in the nucleus. Active domain sizes increase with Pol II cluster radius, control domain sizes do not. h) Same for number of localizations in domains of the respective mark.

h) Cumulative distribution function (cdf) of cluster radius of gyration for chromatin marks and Pol II. Pol II clusters tend to be larger than chromatin clusters. Curves represent mean of cdfs for individual nuclei.

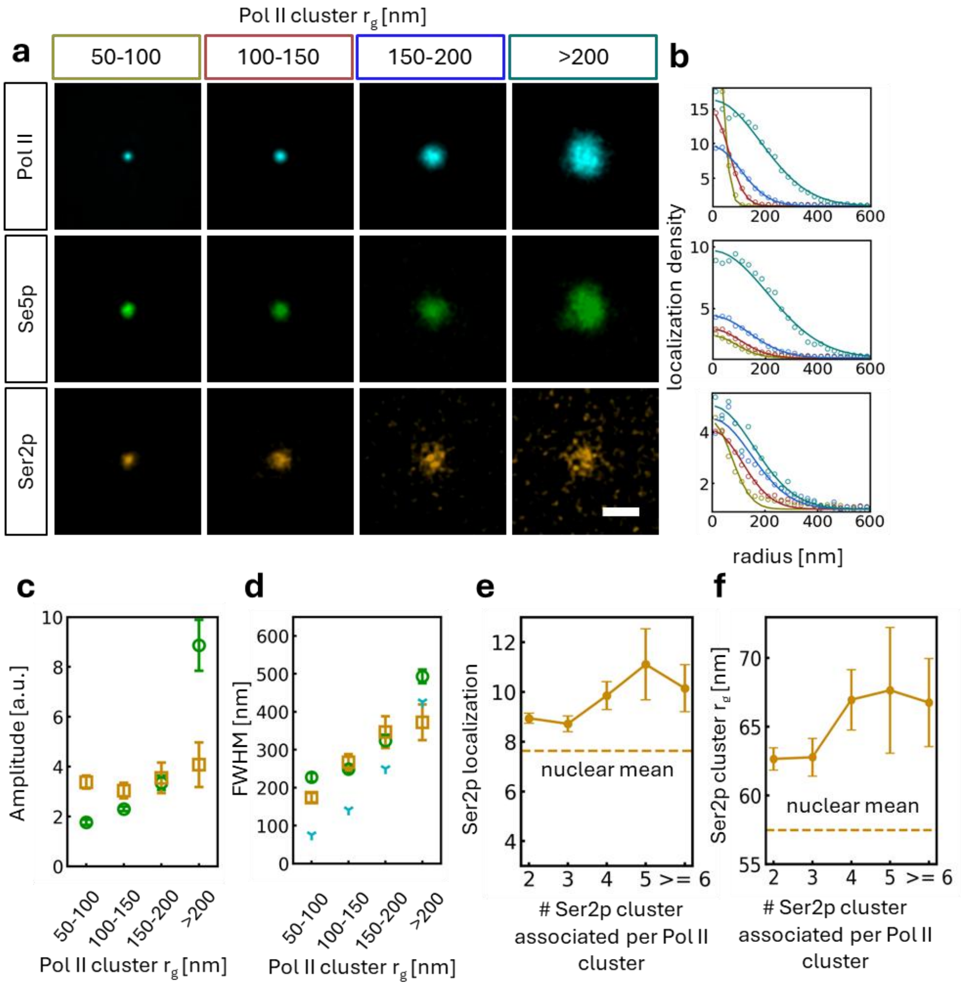

**Suppl. Fig. 6: Pol II clusters enhance transcriptional burst.**

a) Aggregate of Pol II (top), Ser5p (middle), and Ser2p (bottom) localizations aligned by Pol II cluster centroids and grouped by Pol II cluster size class. Scale bar 500nm

b) Normalized localization density (circles) fitted with a Gaussian distribution.

c) Amplitude and d) spread of Pol II phosphor-form localizations obtained from Gaussian fits on b). Mean and standard deviation from bootstrapping (resampling with replacement, 10 iterations). Ser5p density (amplitude) increases strongly with Pol II cluster size.

e) Average size of Ser2p foci in number of localizations and f) radius of gyration as a function of the number of Ser2p foci per Pol II clusters, i.e. the number of co-transcribed elements. Pol II clusters of any size with more transcribed elements result in larger transcriptional bursts. Dashed lines represent the average of all Ser2p foci in the nucleus. Mean and ±s.e.m.

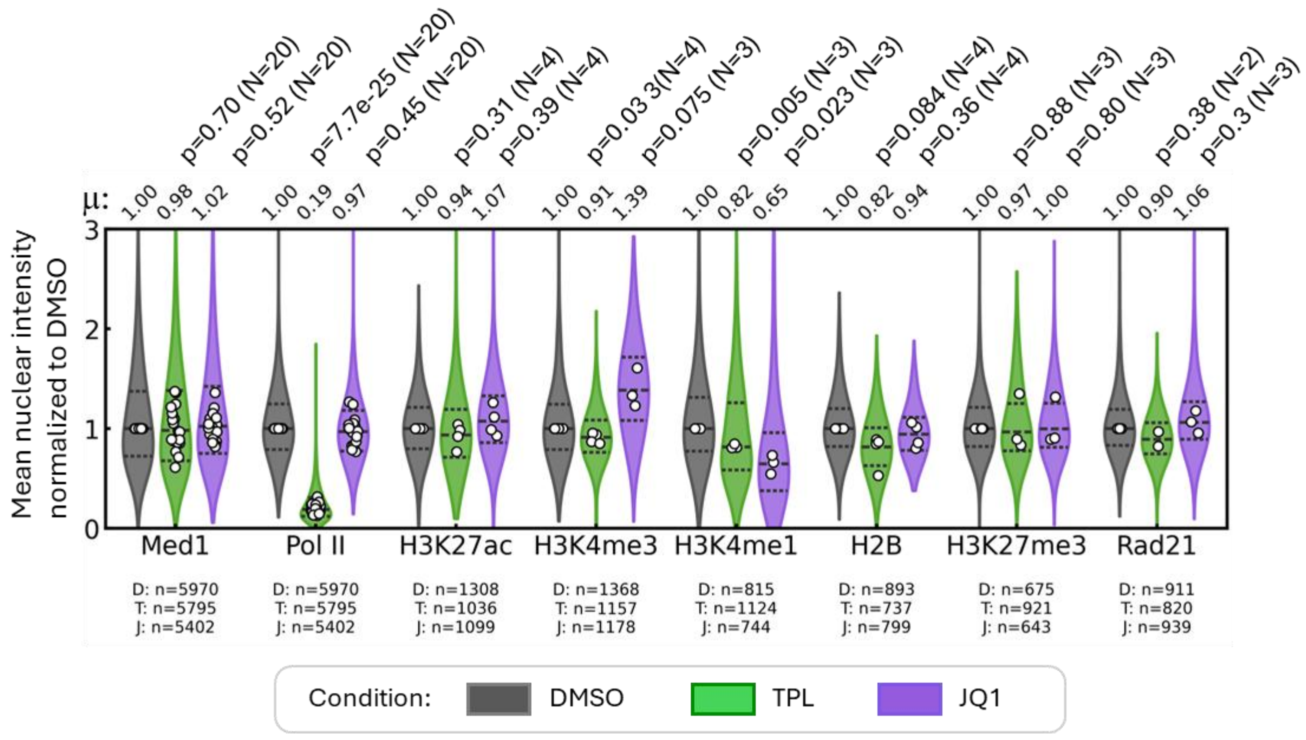

**Suppl. Fig. 7: Absolute signal levels under TPL or JQ1 treatment.**

Violin plots showing the average signal intensity in various channels per individual nuclei under triptolide (TPL) or JQ1 treatment compared to DMSO control. Numbers below the graph refer to the number of nuclei evaluated under DMSO (D), triptolide (T), or JQ1 (J). Values are normalized to the median intensity of the control condition (1.00) and report relative change in the treated conditions.

White circles represent the median value of each of N replicates. The mean (μ) of all replicates for each condition is stated above the graph.

The total number of nuclei (n) throughout the condition is shown below the epitope names.

Paired two-tailed t-test comparing the medians of replicates under TPL or JQ1 to DMSO. p-values (p) along with the number of replicates (N) are shown above the axes.

**Suppl. Table 1:** Antibodies used for immunofluorescence staining in dSTORM microscopy.

| Primary antibodies | | | | | |
|---|---|---|---|---|---|
| **Target (epitope)** | **Source** | **Cat. No.** | **Host species** | **Dilution** | |
| RNA Pol II | Santa Cruz Biotech | sc47701 | Mouse | 1:500 | |
| RNA Pol II | Abcam | ab26721 | Rabbit | 1:500 | |
| Ser5p Pol II | MBL Life Science | Mabi 0603 | Mouse | 1:100 | |
| Ser2p Pol II | Active Motif | 61084 | Rat | 1:100 | |
| H3K27ac | Abcam | ab4729 | Rabbit | 1:500 | |
| H3K4me1 | Active Motif | 39498 | Rabbit | 1:500 | |
| H3K4me1 | Invitrogen | 710795 | Rabbit | 1:500 | |
| H3K4me3 | Active Motif | 39060 | Rabbit | 1:500 | |
| H2B | Abcam | ab1790 | Rabbit | 1:1000 | |
| H2B | Invitrogen | MA5-24697 | Rabbit | 1:1000 | |
| H3K27me3 | Active Motif | 39055 | Rabbit | 1:500 | |
| Rad21 | Abcam | ab217678 | Rabbit | 1:500 | |
| **Secondary antibodies** | | | | | |
| **Target (epitope)** | **Source** | **Cat. No.** | **Host species** | **Dilution** | **Label** |
| Mouse | Abcam | ab150105 | Donkey | 1:500 | AF488 |
| Rabbit | Abcam | ab150075 | Donkey | 1:500 | AF647 |
| Rabbit | Invitrogen | A21206 | Donkey | 1:500 | AF488 |
| Rat | Invitrogen | A78947 | Donkey | 1:500 | AF647 |
| **Antibody combinations** | | | | | |
| | First channel | | Second channel | | |
| Experiment | 1° | 2° | 1° | 2° | |
| Pol II + Ser5p | ab26721 | ab150075 | Mabi 0603 | ab150105 | |
| Pol II + Ser2p | sc47701 | ab150105 | ActiveMotif 61084 | Invitrogen A78947 | |
| Pol II + H3K27ac | sc47701 | ab150105 | ab4729 | ab150075 | |

| | | | | |
|---|---|---|---|---|
| Pol II + H3K4me3 | sc47701 | ab150105 | ActiveMotif 39060 | ab150075 |
| Pol II + H3K4me1 | sc47701 | ab150105 | ActiveMotif 39498 | ab150075 |
| Pol II + H2B | sc47701 | ab150105 | ab1790 | ab150075 (dilution 1:1000) |
| Pol II + H3K27me3 | sc47701 | ab150105 | ActiveMotif 39055 | ab150075 |

**Suppl. Table 2:** Fluorescent probes for 3-color imaging (related to Figure 7).

<table>
<tr><th colspan="2">Direct labeling of condensate constituents</th><th colspan="4">Immunostaining of other protein marks.</th></tr>
<tr><td rowspan="2">First channel</td><td rowspan="2">Second channel</td><td colspan="4">Third channel</td></tr>
<tr><td>epitope</td><td>1°</td><td>2°</td><td>Label</td></tr>
<tr><td rowspan="6">Pol II – Snap-JFX650</td><td rowspan="6">Med1-Halo- TMR</td><td>H3K27ac</td><td>ab4729</td><td rowspan="6">Invitrogen A21206<br><br>(dilution 1:1000 for H2B, everything else 1:500)</td><td rowspan="6">AF488</td></tr>
<tr><td>H3K4me3</td><td>ActiveMotif 39060</td></tr>
<tr><td>H3K4me1</td><td>Invitrogen 710795</td></tr>
<tr><td>H2B</td><td>MA5-24697</td></tr>
<tr><td>H3K27me3</td><td>ActiveMotif 39055</td></tr>
<tr><td>Rad21</td><td>ab217678</td></tr>
</table>
